# Chloroplast Cell-Free Systems from Different Plant Species as a Rapid Prototyping Platform

**DOI:** 10.1101/2024.02.19.580994

**Authors:** Clemens V. Böhm, René Inckemann, Michael Burgis, Jessica Baumann, Cedric K. Brinkmann, Katarzyna E. Lipińska, Sara Gilles, Jonas Freudigmann, Vinca Seiler, Lauren G. Clark, Michael C. Jewett, Lars M. Voll, Henrike Niederholtmeyer

**Author notes:** Corresponding Authors: **René Inckemann** - Max-Planck Institute for Terrestrial Microbiology, Marburg 35043, Germany; Center for Synthetic Microbiology, Philipps-Universität Marburg, Marburg 35032, Germany,; **Lars M. Voll** - Philipps-Universität Marburg, Molecular Plant Physiology, Marburg 35043, Germany; Center for Synthetic Microbiology, Philipps-Universität Marburg, Marburg 35032, Germany,; **Henrike Niederholtmeyer** - Technical University of Munich, Campus Straubing for Biotechnology and Sustainability, Straubing 94315, Germany; Max-Planck Institute for Terrestrial Microbiology, Marburg 35043, Germany; Center for Synthetic Microbiology, Philipps-Universität Marburg, Marburg 35032, Germany. contributed equally (Clemens V. Böhm, René Inckemann).

## Abstract

Climate change poses a significant threat to global agriculture, necessitating innovative solutions. Plant synthetic biology, particularly chloroplast engineering, holds promise as a viable approach to this challenge. Chloroplasts present a variety of advantageous traits for genetic engineering, but the development of genetic tools and genetic part characterization in these organelles is hindered by the lengthy timescales required to generate transplastomic organisms. To address these challenges, we have established a versatile protocol for generating chloroplast-based cell-free gene expression (CFE) systems derived from a diverse range of plant species, including wheat (monocot), spinach, and poplar trees (dicots). We show that these systems work with conventionally used T7 RNA polymerase, as well as the endogenous chloroplast polymerases, allowing for detailed characterization and prototyping of regulatory sequences at both transcription and translation levels. To demonstrate the platform for characterization of promoters and 5’ and 3’ untranslated regions (UTRs) in higher plant chloroplast gene expression, we analyze a collection of 23 5’UTRs, 10 3’UTRs, and 6 chloroplast promoters, assessed their expression in spinach and wheat extracts, and found consistency in expression patterns, suggesting cross-species compatibility. Looking forward, our chloroplast CFE systems open new avenues for plant synthetic biology, offering prototyping tools for both understanding gene expression and developing engineered plants, which could help meet the demands of a changing global climate.

**Graphical Abstract:** 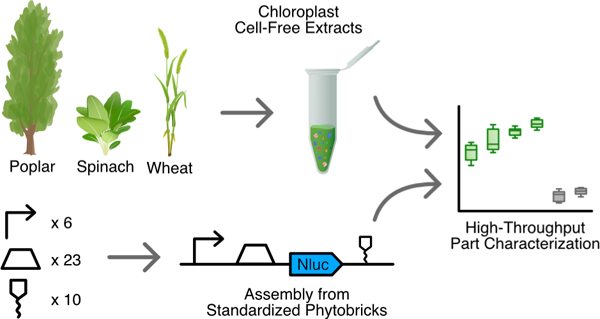

## Introduction

Global agriculture faces a daunting challenge posed by climate change. Rising temperatures, unpredictable weather patterns, and extreme climatic events increasingly threaten crop yields, necessitating the development of more robust agricultural varieties.^1,2^ In this context, plant synthetic biology emerges as a potential solution, offering innovative strategies to enhance the resilience and adaptability of crops to changing environmental conditions.^3,4^ Yet, the current pace of crop engineering is slow, mainly because plant growth is inherently slow. Discovery, development, and approval of novel crop traits currently take about 10 to 15 years, with about 4 years spent developing proof-of-concepts and optimizing genetic constructs.^5^ The genetic system of the chloroplast exhibits several advantageous traits for engineering, such as precise integration of foreign DNA, absence of gene silencing, the option to stack transgenes in synthetic operons, higher predictability of gene expression outcomes and the reduced risk of transgene escape.^6–9^ These features make chloroplasts an attractive target for introducing novel traits into plants.^10,11^

Despite the recognized potential, the field of chloroplast biotechnology is still in its nascent stages. While transplastomic plants have been successfully generated in some species, tools available for chloroplast engineering are still limited.^12^ The foundation of synthetic biology is rooted in the application of engineering principles to biological systems, with standardization being a critical aspect.^13^ Standardization is particularly vital for the development and dissemination of easily shareable biological parts.^14^ The availability of a diverse array of genetic parts for the controlled expression of transgenes is especially essential in plastids. This need arises from the potential risks associated with the repeated use of identical genetic elements in the chloroplast genome, which could lead to unintended homologous recombination, even with sequence stretches as short as 50 base pairs^15^ Moreover, synthetic genetic circuits often require the precise calibration of their constituent parts’ activities to function reliably.^16,17^ To enable synthetic biology applications in plastids to the level of versatility and complexity already achieved in bacteria would require the availability of well-characterized gene expression elements with a broad range of activities.

However, genetic parts characterization in plastids is more challenging than in microbes due to longer timescales needed for generating transplastomic organisms. Extensive and efficient genetic tools enable rapid engineering within days in microbial chassis. In contrast, obtaining homoplasmic plastid transformants and testing parts in vivo requires several months of selection, or in case of poplar trees even up to an entire year.^18–20^ In order to overcome these limitations, rapid prototyping platforms that facilitate an accelerated Design-Build-Test-Learn cycle (DBTL) need to be developed.^21^ Such a platform would enable rapid iterations of genetic designs, testing, and modifications, significantly reducing the time and resources required for part characterization in plastids.

Cell-free gene expression (CFE) systems^22–24^, which have already accelerated engineering and characterization of genetic parts in non-model bacteria^22–26^, yeast^27^ and mammalian cells^28^, present a viable solution for rapid prototyping in chloroplast biotechnology. These systems allow for the in vitro analysis and testing of genetic components, bypassing the need for whole-plant transformations. Notably, chloroplast lysates have a history of use in studying gene expression regulation that led to foundational discoveries in chloroplast biology, such as the regulation of transcription and translation by light, the effect of regulatory nuclear proteins and other environmental stimuli.^29–36^

In this study, we aim to investigate the feasibility of cell-free prototyping using chloroplast extracts from a diverse range of plant species, encompassing both agricultural crops and tree species. We focussed on testing regulatory components involved in post-transcriptional and translation control, as these stages are the primary ways for the regulation of gene expression in chloroplasts^37^. Our work builds off a recently established protocol for generating a highly active tobacco chloroplast-based cell-free transcription and translation system, which demonstrated the feasibility of using CFE systems for characterization of plant-based ribosome binding sites.^38^

We report the development of chloroplast-based CFE systems derived from wheat (monocot), spinach, and poplar trees (dicots). These chloroplast crude extracts were obtained by isolating intact plastids using density gradients, lysing them, and then using ultracentrifugation to remove cell debris and retain key components of the protein biosynthesis machinery, enabling in vitro transcription and translation. We develop an automated workflow to facilitate high-throughput part characterization while conserving valuable reagents. By using this workflow we conducted a comprehensive characterization of 38 distinct genetic elements, originating from various plant species, bacteria, viruses, and synthetic sources, encompassing 5’ untranslated regions (UTRs), 3’UTRs, and endogenous chloroplast promoters. Our results demonstrate part transferability across different plant species. Given the slow growth and challenging engineering of many plant species, we anticipate that rapid cell-free testing will accelerate plant synthetic biology.

## Result and Discussion

### Development of High-Yielding Cell-Free Expression Systems from Wheat, Spinach, and Poplar

Our objective was to develop a high-yielding CFE system, derived from wheat (a monocot), as well as spinach and poplar trees (dicots). We adapted the workflow for generating highly active cell-free transcription and translation systems from tobacco chloroplasts (Supplementary Figure S1). For the harvesting process, plant leaves were cut into smaller pieces and homogenized. Subsequently, a centrifugation step separated chloroplasts from the rest of the leaf material. This was followed by gradient centrifugation, employing a stepwise Percoll gradient to separate intact chloroplasts from the broken ones.

To prepare cell-free systems, the contents of the intact chloroplasts were released by disrupting the envelope membranes. For lysis, chloroplasts were passed through 25G needles, with the number of passes (15-40 times) varying depending on the species. For each isolation, approximately 100–300 grams of leaves were harvested, yielding about 1 ml of spinach or 200 µl of wheat or poplar cell-free extract. The resulting extracts could be stored at -80°C at least for two years without significant loss of activity. Although refreezing in liquid nitrogen after thawing was possible, the activity declined over multiple freeze-thaw cycles.

Adapting our CFE systems to a variety of plant species required subtle modifications at multiple workflow stages. Key among these was addressing the unique growth conditions and harvest timing for each species. Additionally, the distinct physical properties of each plant’s tissue necessitated tailored homogenization methods to ensure consistent processing. Due to variability in chloroplast size, volume, and density between species, we had to adapt the stepwise density gradients to each species to ensure successful chloroplast isolations (see Methods).

In the subsequent cell-free reactions, we combined chloroplast cell-free extracts with various DNA constructs. Here, we established an automation workflow, leveraging an Echo 525 liquid handler for precise acoustic liquid handling (Figure 1). This technology has been shown previously in cell-free systems^24,28,39,40^ and was pivotal for downsizing the reaction volumes to conserve extract volume and to increase the number of testable constructs. In conjunction with this, we utilized a contactless nanoliter dispenser to set up the luciferase assays. This automation approach not only streamlined the process but also enabled a significant increase in the throughput of part characterizations (Figure 1).

**Figure 1:**
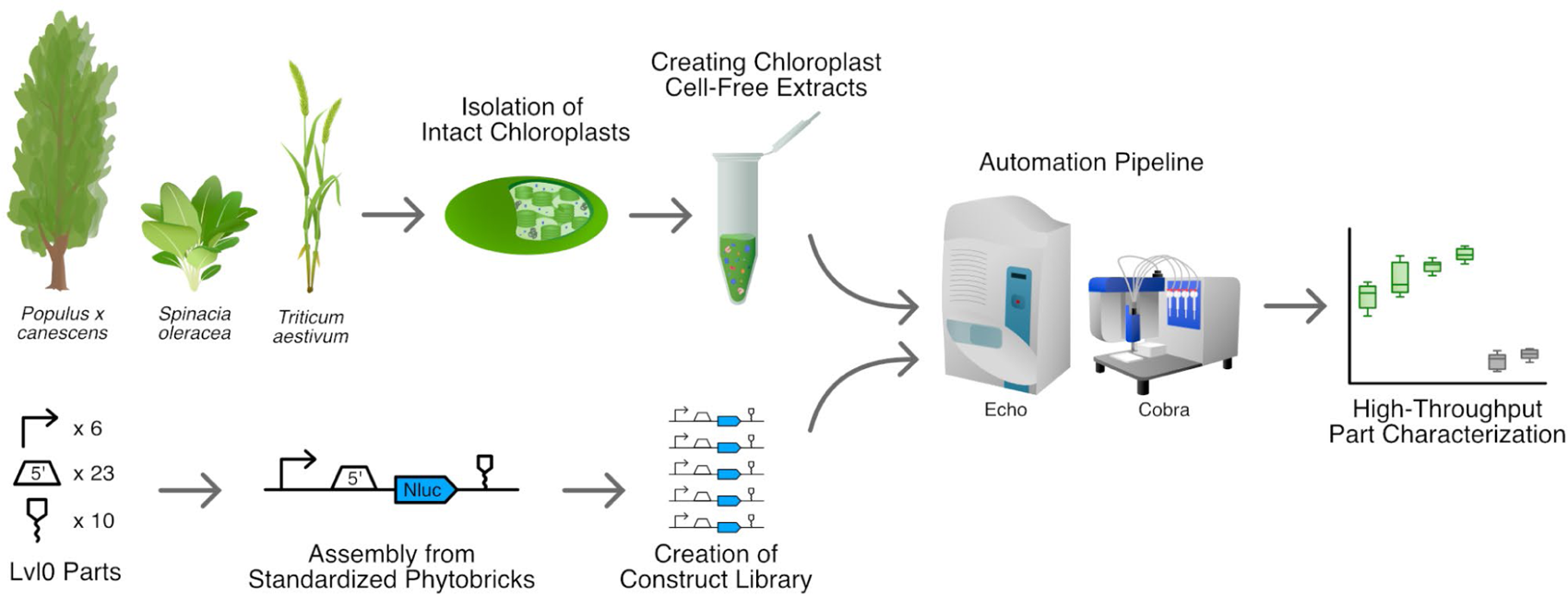
Workflow for employing chloroplast cell-free systems from various plant species (poplar, wheat, spinach) for automated high-throughput part characterization. Chloroplast cell-free extracts were generated from *Populus × canescens* (poplar), *Spinacia oleracea* (spinach) and *Triticum aestivum* (wheat) via the isolation of intact chloroplasts and subsequent lysis. A library of standardized Phytobrick Level 1^14^ assemblies was then constructed and tested, comprising diverse regulatory elements. Cell-free reactions were set up via an automated workflow involving contactless liquid handlers (Echo 525, Cobra) to combine chloroplast cell-free extracts with DNA templates and NanoLuc substrate.

### Demonstrating Translation Activity of the Chloroplast Cell-Free Extracts

We first aimed to validate whether the chloroplast CFE systems possessed sufficient translation activity for characterization of parts. To this end, we employed the NanoLuc reporter system, noted for its high sensitivity and recent application in cell-free systems.^41^ To facilitate our initial experiments, we engineered an ‘universal test construct’, designed to be a standard tool for troubleshooting chloroplast cell-free extracts. This construct comprised genetic elements of viral origins, specifically the T7 RNA polymerase promoter, gene10 5’UTR, and the Tobacco Mosaic Virus (TMV) 3’UTR, each chosen for their proven efficacy in driving strong gene expression in chloroplasts. To support broader research efforts, we have made this ‘universal test construct’ accessible to the scientific community through Addgene (ID 216625). To evaluate the capacity for cell-free transcription and translation, we combined the chloroplast extract with a reaction buffer and the DNA template, then incubated for 4 hours for the NanoLuc reporter to accumulate. To assess synthesis of the reporter, we combined the reaction mixture with the NanoLuc substrate for Luminescence measurements.

Throughout these experiments, transcription was facilitated using supplemented T7 RNA polymerase, allowing us to focus on verifying the creation of translationally active extracts. We successfully detected NanoLuc luciferase signals from spinach, wheat, and poplar extracts. Notably, the luminescence signals observed were more than 1000-fold greater than the background signal from the ‘no extract’ negative control (Figure 2A).

**Figure 2:**
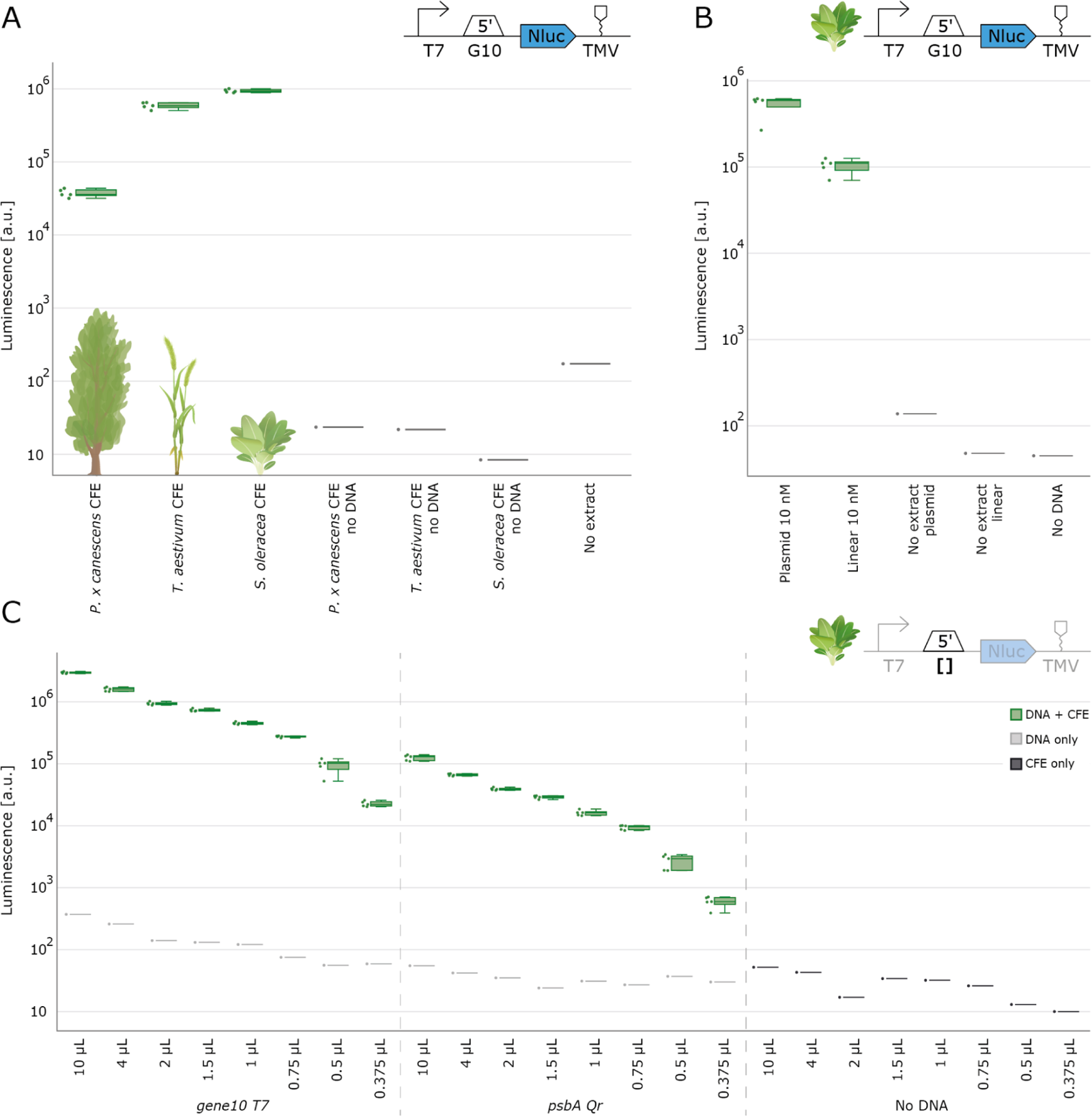
Validation of cell-free protein synthesis by chloroplast extracts from different plant species. (a) NanoLuc luminescence signal for cell-free reactions from poplar, spinach and wheat, including negative controls without extract or without DNA template (b) Comparison of NanoLuc luminescence signal from cell-free reactions with plasmid and linear DNA templates (c) NanoLuc luminescence signal from cell-free reactions with different final reaction volumes to demonstrate downscaling potential. In (a) and (b), cell-free reactions were set up with a total volume of 2µl. NanoLuc activity was measured after 4 hours of incubation at 20°C (N=5).

Utilizing purified NanoLuc protein for comparison, we demonstrated that our cell-free reactions yielded NanoLuc protein at levels up to 20 nM (Supplementary Figure S2). While the signal strengths for spinach and wheat were comparable, the luminescence from poplar extracts was 20 times lower. However, the signal from poplar still exceeded the background of the negative control by approximately 100-fold. Based on these results, we deemed the poplar extracts sufficiently functional for further part characterization. Optimizing the extraction process or adjusting the cell-free reaction mixture could enhance the performance of the poplar extracts in the future. Such improvements will potentially elevate the protein synthesis efficiency to levels more consistent with those observed in spinach and wheat extracts.

We also assessed the feasibility of employing linear DNA templates, without the implementation of any specific optimizations or adjustments to the cell-free reaction mixture.We observed that linear DNA templates produced substantial luminescence signals, indicating successful expression (Figure 2c). However, the signal intensity from linear templates was almost one order of magnitude lower compared to that obtained from plasmid DNA. Degradation of linear DNA or other molecular interactions within the system could have caused this reduced efficiency. These findings suggest that, upon further optimization, linear DNA templates may be effectively used for part characterization. Such an approach offers several advantages, with the most prominent being the bypass of time-consuming, in vivo cloning procedures. We decided to use plasmid DNA in our current system to more reliably characterize genetic elements that confer a wide range of expression strengths, as parts resulting in low expression would be more difficult to measure using linear DNA.

To identify the most effective DNA concentration for our characterization experiments, we carried out a DNA concentration titration (Supplementary Figures S3 and S4). Notably, expression levels plateaued at a concentration of 10 nM DNA, which we subsequently used for all following experiments. To evaluate a broader range of genetic parts, we established an automated setup to systematically reduce and optimize the reaction volume of our cell-free system. Our findings revealed that the reaction volume could be effectively minimized to 375 nl, still yielding a NanoLuc signal 300-fold over the negative control’s background using the universal test construct (Figure 2c). Nonetheless, we opted for a final volume of 2µl, considering that a lower expressing construct employing a *psbA* 5’ UTR showed expression only 20-fold over the background at this low volume, and we aimed to ensure effective characterization of parts with low expression levels.

### Quantitative Characterization of 5’UTRs for Chloroplast Expression

After confirming the functionality of our extracts and establishing a high-throughput automated workflow, we next sought to determine the suitability of chloroplast cell-free extracts for the systematic and quantitative characterization of genetic components in chloroplast gene expression. To this end, we constructed a genetic part library comprising 23 distinct constructs, differing in their 5’UTR sequences. The 5’ UTR is recognized as a key player in gene expression regulation, due to its roles in initiating translation and harboring mRNA stabilizing elements, which prevent mRNA degradation. The 5’UTRs were obtained from various origins, including chloroplast genomes of diverse plant species such as wheat (Ta), oak (Qr), tobacco (Nt), rice (Os), and spinach (So), *Escherichia coli* phage T7, and synthetic sources (including Ribosome Binding Sites from the iGEM registry, originally designed for *E. coli*). Except for the 5’UTR, we kept all other sequence elements constant, employing the T7 promoter and the TMV 3’UTR consistent with the configuration of the universal test construct.

We tested all constructs in a spinach chloroplast cell-free extract, with luminescence measured after 4 hours of incubation. NanoLuc signal was detected for all constructs and was at least an order of magnitude higher than the ‘no DNA’ and ‘no extract’ controls (Figure 3). We utilized a ‘dummy’ part, lacking elements for mRNA stabilization and initiation of translation as an additional negative control. Generally, in our experiments, the ‘no extract’ control exhibited a higher signal compared to the ‘no DNA’ control, primarily because of the slight co-purification of luciferase protein from *E. coli* during plasmid preparation, which leads to luminescence due to the high sensitivity of the NanoLuc system. This phenomenon is even more pronounced when using endogenous chloroplast promoters, as they also induce high gene expression in *E. coli*. We found that boiling plasmid DNA for 30 min effectively removes background NanoLuc signals, when needed. For all DNA constructs depicted in Figure 3, NanoLuc signals from the ‘no extract’ controls were 10 to 10,000 times lower than the actual signals obtained using extracts (Supplementary Figure S5).

**Figure 3:**
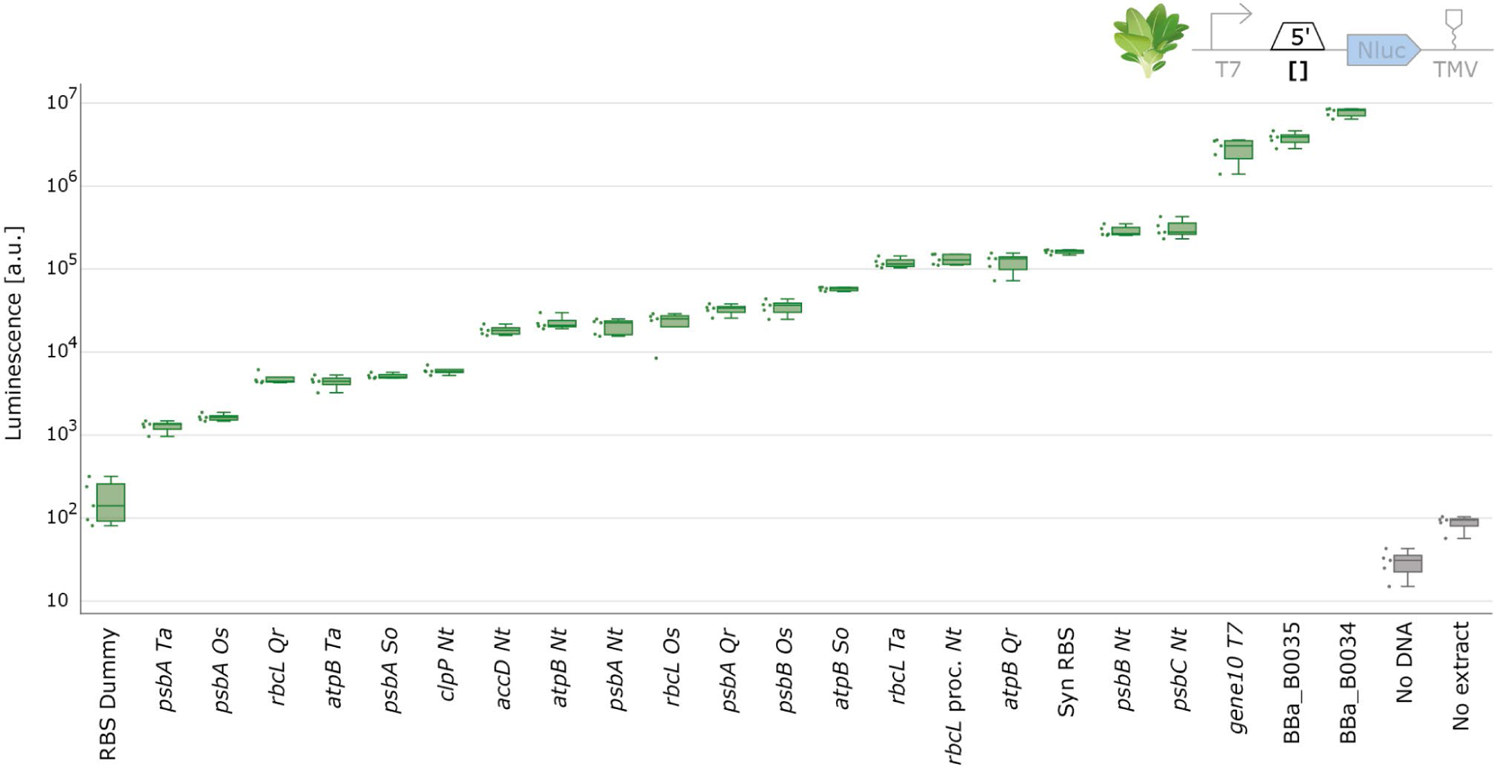
5’UTR characterization with spinach chloroplast cell-free extract. NanoLuc luminescence signals obtained with DNA templates with varying 5’UTR. Negative controls either lack extract or DNA. Cell-free reactions were set up with a total volume of 2µl and NanoLuc activity was measured after 4 hours of incubation at 20°C (N=5).

The data from the 5’UTRs exhibited a broad spectrum of NanoLuc luminescence levels, indicating different translation efficiencies conferred by the 5’UTRs. Our results align with the known importance of 5’ UTRs in regulating gene expression levels, as in plastids expression is mainly controlled on a translational level, due to the 5’UTRs’ involvement in initiating translation and containing mRNA stabilizing elements.^42^ Consistent with findings from in vivo experiments, our data showed that the gene10 5’UTR yielded higher expression levels than the *rbcL* 5’UTR from tobacco.^43^ Furthermore, the ‘RBS dummy’ part displayed the lowest expression, as expected, due to its absence of critical elements for mRNA stabilization and initiation of translation. The high expression strength observed in the BBa_B0034 and BBa_B0034 RBS parts can be attributed to these ribosome binding sites closely resembling the consensus chloroplast Shine-Dalgarno Sequence, with only a single base pair difference.^44^

Contrary to expectations from in vivo studies, our results revealed a lower expression strength for all of the *psbA* 5’UTRs, where a much higher expression would typically be anticipated. This discrepancy could be attributed to the absence of regulatory factors that are usually imported from the nucleus, particularly since *psbA* is known for its complex regulation of translation initiation, involving various RNA binding proteins.^45–47^ A potential next step to address this could involve enhancing the expression strength of the *psbA* 5’UTR by supplementing our system with these regulatory nuclear proteins. Interestingly, parts derived from different plant species still generated a NanoLuc signal in spinach CFE systems, suggesting the potential cross-species utility of these parts in chloroplast engineering.

### Analysis of 3’UTRs in Chloroplast Gene Expression

Following the characterization of 5’UTRs, our next objective was to systematically characterize 3’UTRs, employing a similar approach. We developed a library of 10 unique constructs, each distinguished by its 3’UTR sequence, and tested them in spinach CFE. As anticipated from in vivo studies, variations in 3’UTR only had small effects on expression compared to 5’UTRs. Yet there was still a notable variation, approximately one order of magnitude difference, between the lowest and highest expressing constructs (Figure 4).

**Figure 4:**
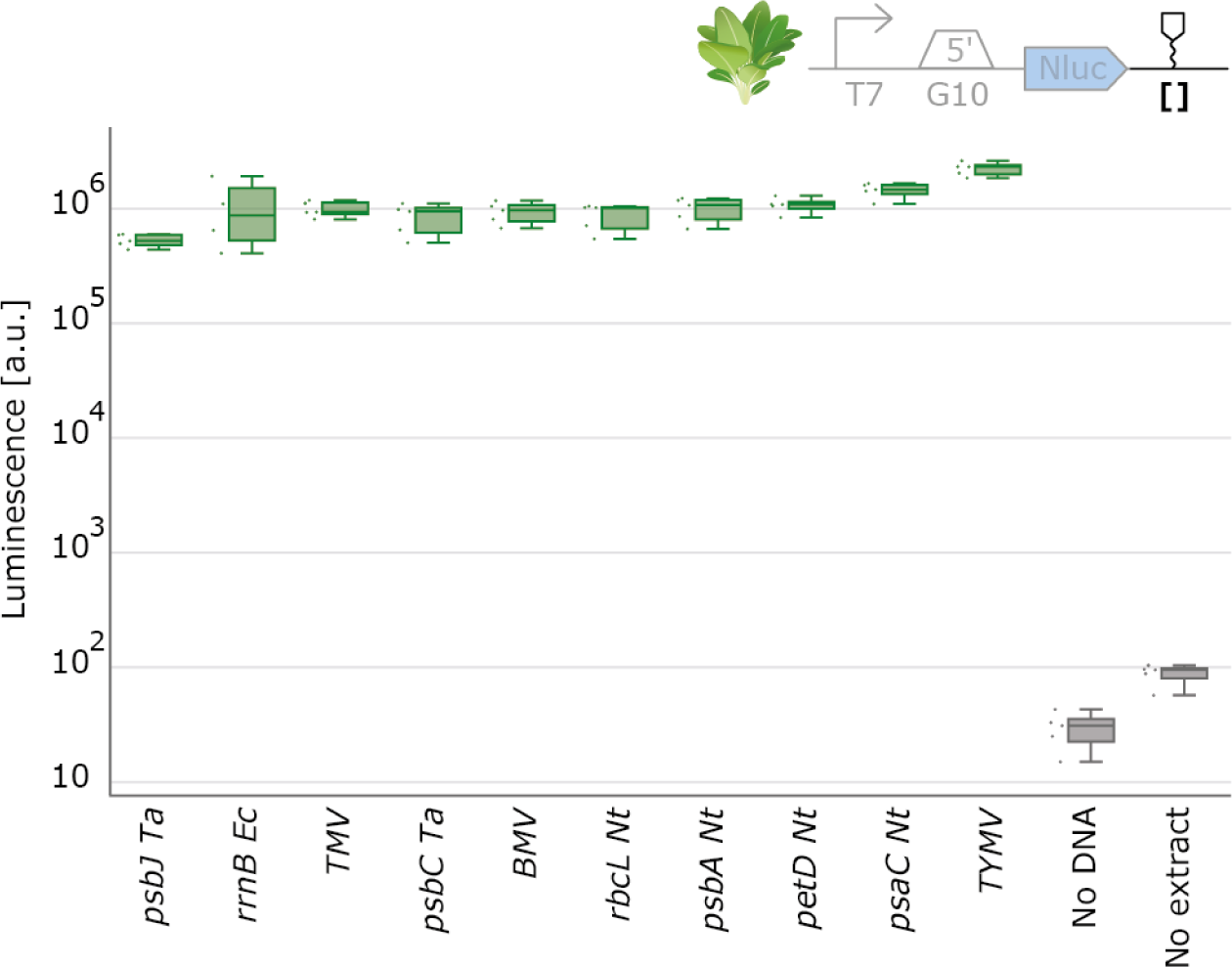
3’UTR characterization with spinach chloroplast cell-free extract. NanoLuc luminescence signals obtained with DNA templates with varying 3’UTR. Negative controls either lack extract or DNA. Cell-free reactions were set up with a total volume of 2µl and NanoLuc activity was measured after 4 hours of incubation at 20°C (N=5).

Aligning with the conclusions of previous in vivo studies, our data reveals that 3’UTRs hold a relatively minor role in determining final protein levels in chloroplasts.^42,48^ Nevertheless, 3’ UTRs could play a role in fine-tuning expression levels, particularly for proteins such as hetero multimers that require expression at subtly varied levels. Our observations for the 3’UTR characterization again indicate that genetic parts from various plant species will be transferable and functional across different plant species, showcasing their versatility in chloroplast synthetic biology applications.

### Characterization of Genetic Parts in Monocot Crop Species

We next aimed to extend the characterization of the same genetic parts above to wheat, a monocot crop species. This step was crucial to understand how these parts behave in different plant species, given the evolutionary distance between them. For the sake of comparability, we employed the identical constructs with wheat chloroplast cell-free extracts instead of spinach. This direct comparison allowed us to assess the performance of the genetic components in the physiological environment of a distant species.

**Figure 5:**
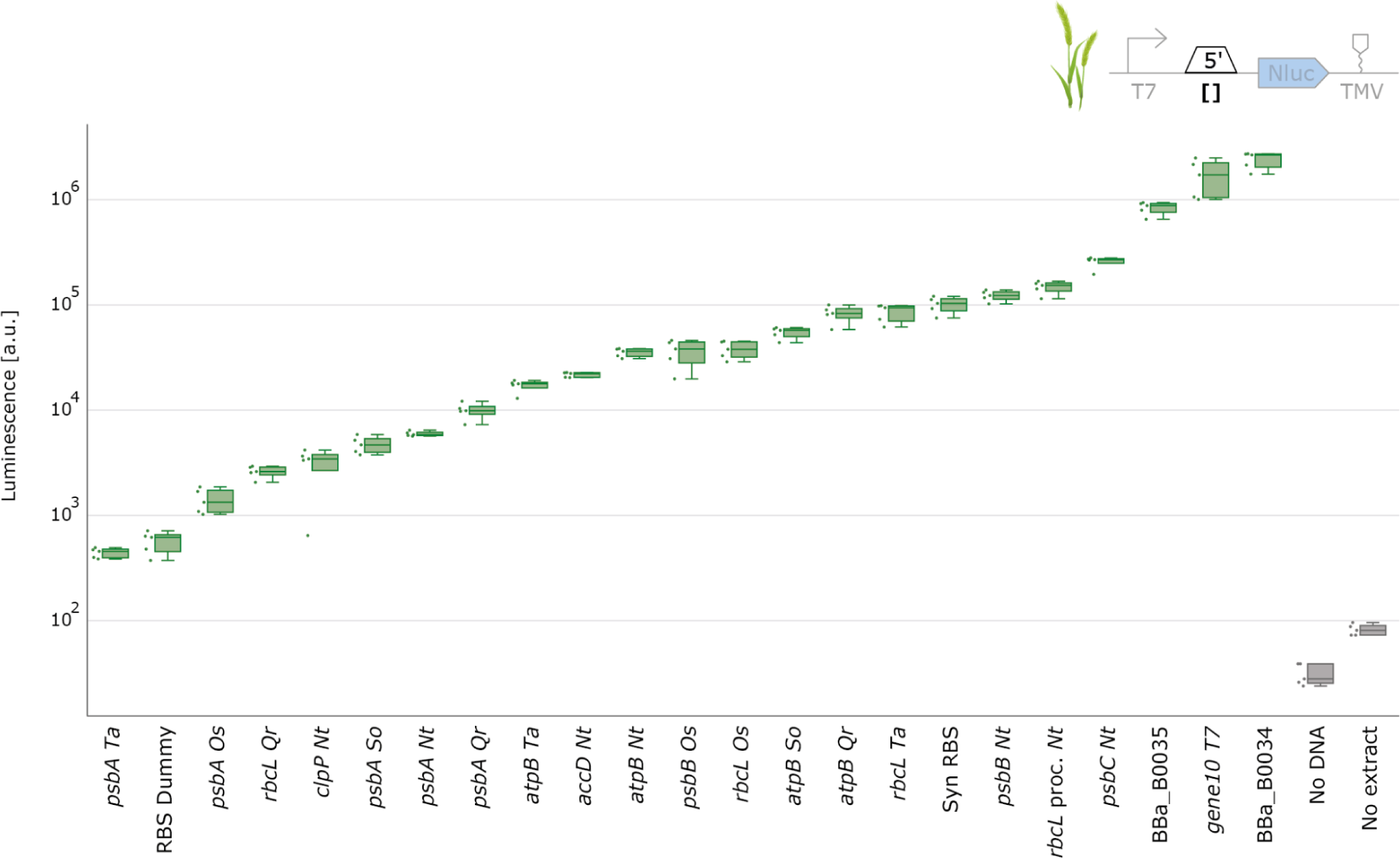
5’UTR characterization with wheat chloroplast cell-free extract. NanoLuc luminescence signals obtained with DNA templates with varying 5’UTR. Negative controls either lack extract or DNA. Cell-free reactions were set up with a total volume of 2µl and NanoLuc activity was measured after 4 hours of incubation at 20°C (N=5).

**Figure 6:**
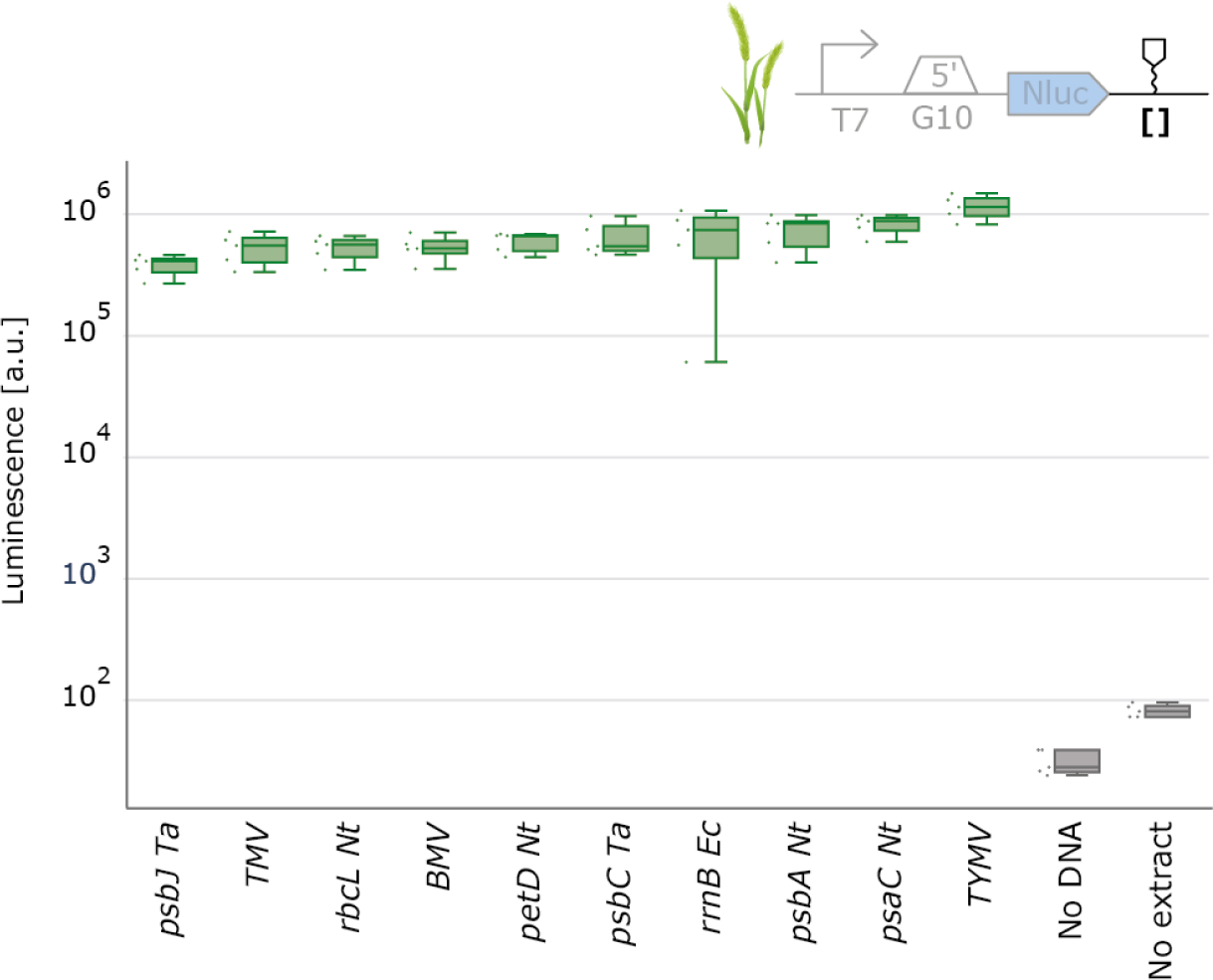
3’UTR characterization with wheat chloroplast cell-free extract. NanoLuc luminescence signals obtained with DNA templates with varying 3’UTR. Negative controls either lack extract or DNA. Cell-free reactions were set up with a total volume of 2µl and NanoLuc activity was measured after 4 hours of incubation at 20°C (N=5).

Our results showed a wide range of expression strengths for the various 5’ UTRs in wheat (Figure 5), mirroring the trends observed in spinach. Similarly, the 3’ UTRs in wheat also exhibited a comparable range of expression as seen in spinach (Figure 6). Taken together, our findings underscore the potential of certain genetic parts to function effectively across different plant species, even those as evolutionarily distant as monocots and dicots. Intriguingly, despite the evolutionary divergence between these two plant species, the data from wheat closely aligned with that obtained from spinach.

### Correlation Analysis of Part Performance in Spinach and Wheat

To gain a deeper insight into the transferability of genetic parts, we undertook a comparative analysis between spinach and wheat, focusing on genetic part characterization data. Therefore, we performed a regression analysis via linear regression on log-log transformed data (Figure 7).

**Figure 7:**
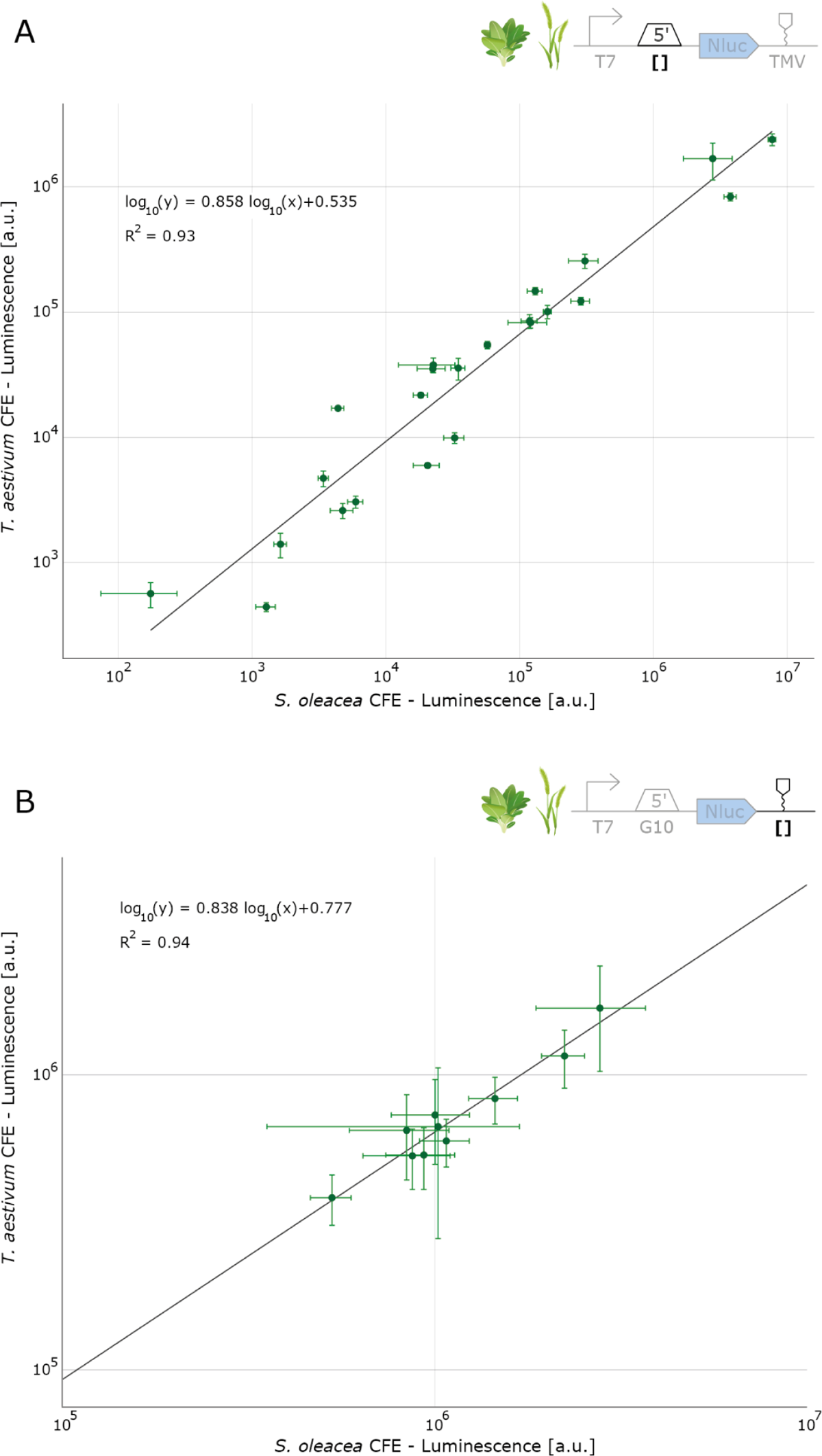
Correlation of UTR performance between spinach (*Spinacia oleracea*) and wheat (*Triticum aestivum*) cell-free chloroplast extracts. (a) Correlation of 5’UTRs performance (b) Correlation of 3’UTRs performance. Regression analysis was performed via linear-least squares regression on log-log transformed data. Each data point represents the mean of five replicates (N=5), the error bars depict the standard deviations.

Our findings revealed a significant correlation in the expression levels of individual constructs between spinach and wheat, with an R^2^ value of 0.93, in case of the 5’UTRs and 0.94 for the 3’UTR performance, indicating a strong relationship. This high similarity in expression across these species, despite their evolutionary distance, highlights a key advantage of chloroplast genome engineering. The relative conservation of chloroplast genomes compared to nuclear genomes could explain this cross-species compatibility.

However, our analysis also identified certain outliers, underscoring the importance of developing chloroplast cell-free extracts in each species of interest, rather than relying solely on one single system. Among the outliers is the *atpB* 5’UTR, derived from the wheat chloroplast genome. Notably, the NanoLuc activity for this 5’UTR in wheat extract was fourfold higher than in spinach extract, hinting at enhanced performance of the *atpB* 5’UTR within its native species context. This observation aligns with existing literature, which has demonstrated that various factors are crucial for translating plastid *atpB* mRNA, particularly due to the absence of a Shine-Dalgarno sequence. These elements include mRNA binding proteins, which may vary between different plant species.^49^

For the other 5’UTRs we tested, this specific phenomenon was less pronounced, underscoring the necessity for a broader comparative analysis of 5’UTRs among diverse plant species. Understanding these nuances could be invaluable in deciphering the mechanisms of translation regulation in the chloroplasts of different plant species.The observed outliers emphasize the variability that can occur due to species-specific genetic and physiological differences. However, our results suggest that spinach chloroplast cell-free extracts might potentially be used to predict part performance in other chloroplast CFE systems during early optimization stages. This approach could bypass the time-consuming steps of growing specific plant species for initial tests, leveraging the ready availability of spinach leaves, also for labs that do not specialize in plant biology. Such a strategy could significantly expedite the preliminary phases of chloroplast genetic engineering, particularly in species where growth conditions and development times are limiting factors.

### Establishing an Endogenous Chloroplast Transcription/Translation System

In all previous experiments, we utilized T7 RNA polymerase to drive transcription. To test if besides translation, the transcription machinery of our chloroplast extracts was active, we selected our previously highest expressing 5’UTR (BBa_0034) and 3’UTR (TMV), which we combined with putatively strong native chloroplast promoters, such as the P*_rrn16_*, P*_rbcL_* or P*_psbA_* promoters. The experiment involved testing five distinct promoters, maintaining all other regulatory elements constant. Additionally, we incorporated a positive control that relied on T7 polymerase for comparative purposes. Notably, we were able to detect a NanoLuc signal in this setup, indicating the successful reconstitution of a completely endogenous chloroplast transcription/translation system (Figure 8).

**Figure 8:**
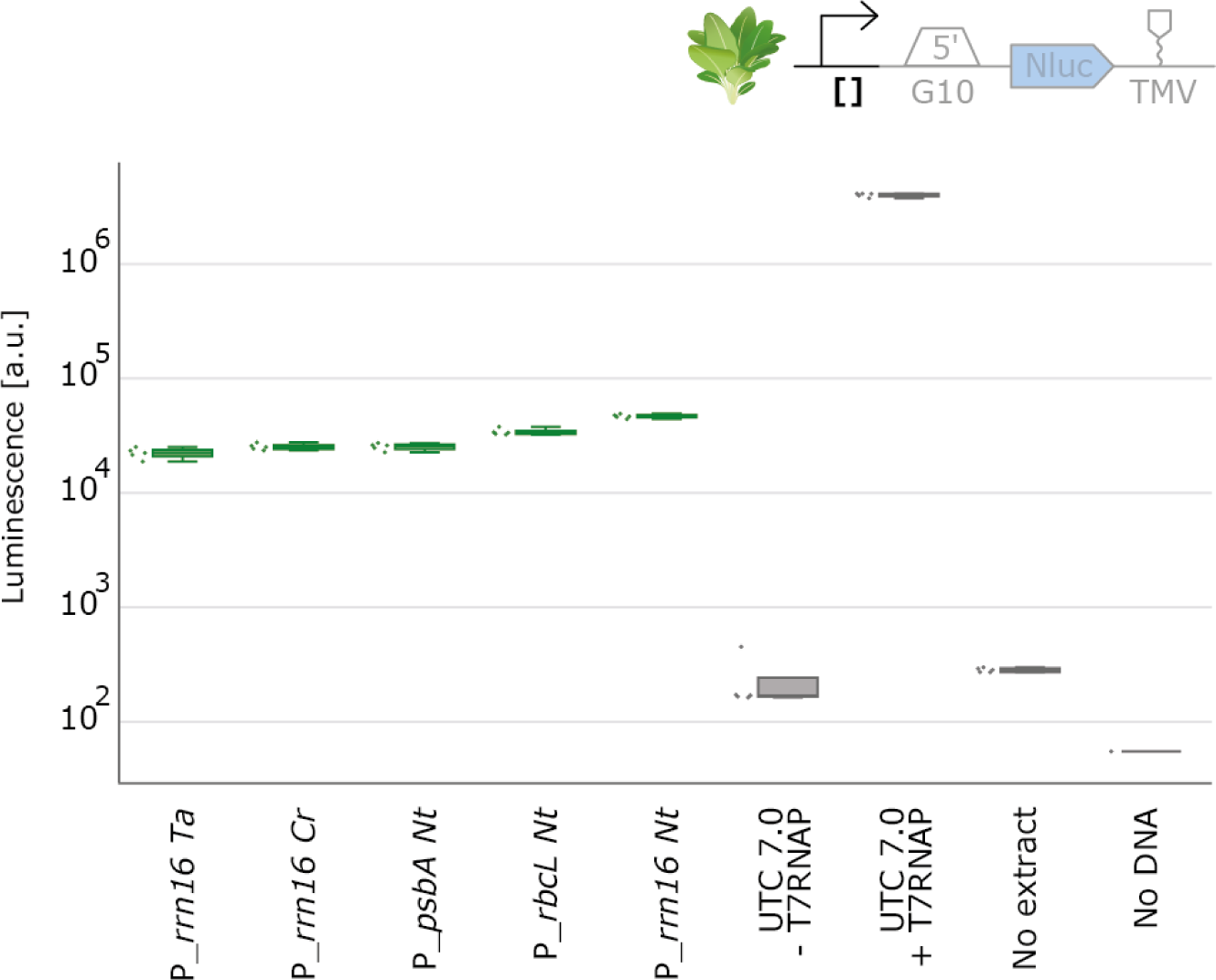
Endogenous transcription in chloroplast CFE reactions. NanoLuc luminescence signals obtained with DNA templates with varying 5’UTR. Negative controls either lack extract or DNA. Cell-free reactions were set up with a total volume of 10µl and NanoLuc activity was measured after 6 hours of incubation at 20°C (N=5).

This outcome validates the functionality of our endogenously driven system, which relies on the transcription activity of the endogenous chloroplast polymerases. Of note, the signal intensity was approximately 100 times lower compared to the system using T7-based transcription, which may limit our system to characterizations of strong promoters until further optimization.

### Characterization of Genetic Parts in Poplar - A Dicot Tree Species

As the final step in our part characterization study, we focused on poplar, a dicot tree species. Given the poplar extract’s lower protein production capabilities (Figure 2), we aimed to assess whether poplar chloroplast CFE can be effectively utilized for part characterization. For this experiment, we selected a limited set of three genetic parts. To compensate for the anticipated lower protein yield and to enhance the overall signal in NanoLuc measurements, we increased the total reaction volume to 10µl. Remarkably, we found the data from poplar to be comparable to those of spinach and wheat in terms of relative part performance, with RBS 34 producing the highest luminescence values compared to the other 5’UTRs tested and *rbcL* the lowest (Figure 9).

**Figure 9:**
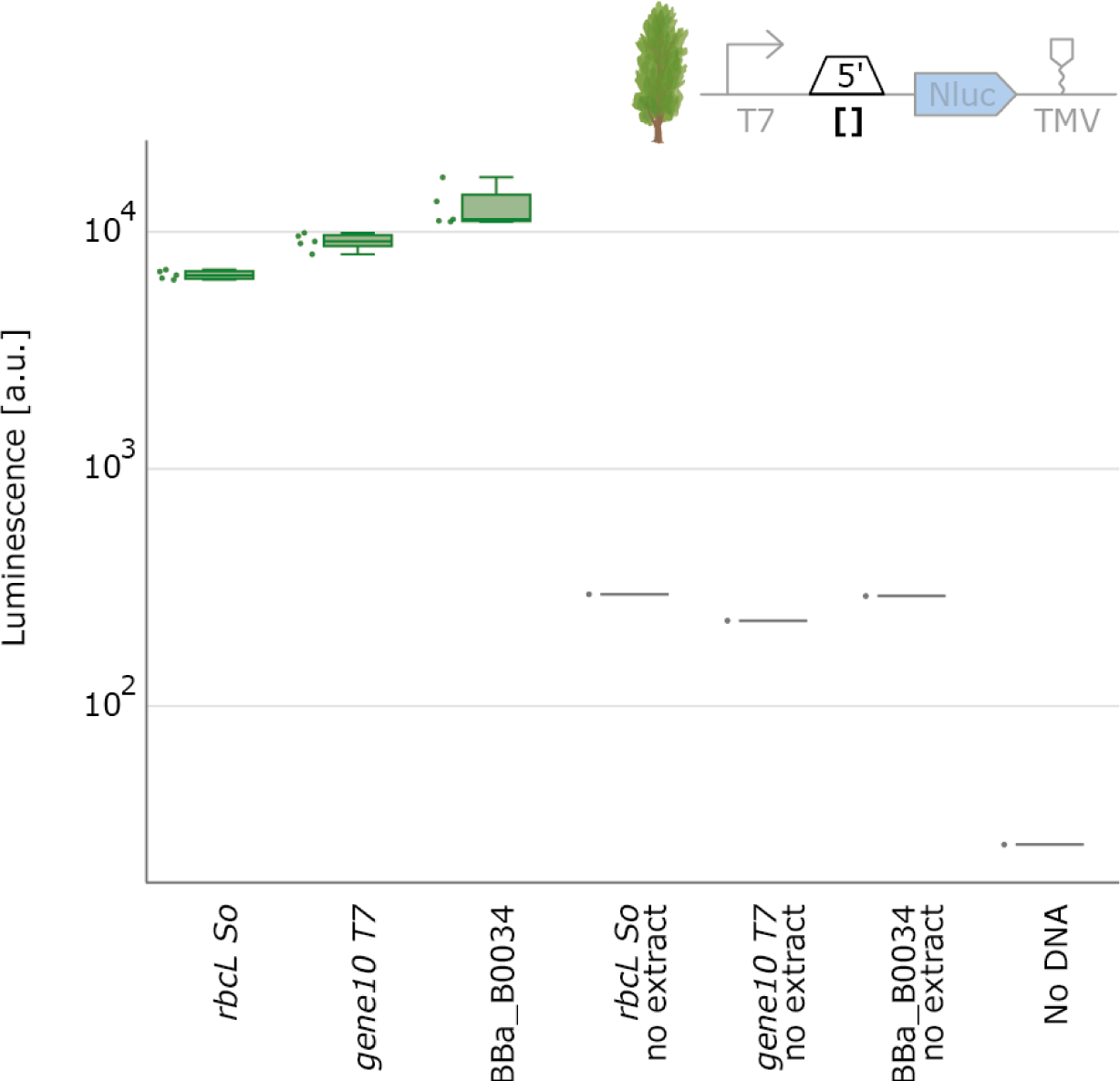
5’UTR characterization with poplar chloroplast cell-free extracts. NanoLuc luminescence signals obtained with DNA templates with varying 5’UTR. Negative controls either lack extract or DNA. Cell-free reactions containing T7 RNA polymerase were set up with a total volume of 10µl and NanoLuc activity was measured after 6 hours of incubation at 20°C (N=5).

The successful characterization of parts in poplar highlights a significant advantage of chloroplast cell-free systems. In vivo part characterization in trees like poplar may take years due to their slower growth and longer generation time. In contrast, chloroplast cell-free systems offer the potential to substantially accelerate the DBTL cycle for chloroplast engineering in trees. By enabling the initial testing of parts in vitro, only the final iterations of a construct would need to undergo the more time-consuming process of in vivo chloroplast transformation. This approach could greatly expedite the development and optimization of genetic modifications in tree species.

## Conclusion

Our study successfully demonstrated the development of CFE systems from the chloroplasts of spinach, wheat, and poplar, demonstrating that a previously developed protocol for tobacco^38^ is versatile across species. We also showcased the potential for native transcription and the characterization and prototyping of regulatory sequences at both transcription and translation levels. These chloroplast cell-free systems provide a powerful platform for the extensive screening of DNA sequences, facilitating high-throughput part characterization. Large-scale characterization of 5’ and 3’UTRs in chloroplast gene expression for higher plants represents a useful resource for future chloroplast engineering efforts.

The next step would need to involve conducting systematic comparisons between cell-free results and in vivo expression to understand the limitations of these chloroplast cell-free systems in prototyping. However, based on a small number of parts that in vivo data exists for, chloroplast extracts produced the expected results. An intriguing aspect to explore is the impact of nuclear factors, which regulate gene expression in chloroplasts in vivo, but might be absent in our cell-free extracts. These factors could potentially be purified individually and incorporated into the cell-free systems.

In summary, the chloroplast cell-free systems we have established are poised to make several impacts. First, as has been done before^29^, they can help elucidate fundamental aspects of chloroplast biology, such as regulation of transcription and translation, which is challenging to study in vivo, especially for essential genes. Second, chloroplast cell-free systems hold promise for expediting the development of novel transplastomic crop varieties, potentially playing a significant role in adapting plants to climate change and enhancing yields through engineered carbon fixation and photosynthetic light reactions.

## Materials and Methods

### Plant growth for chloroplast isolation

*Triticum aestivum* (wheat) and *Populus × canescens* (poplar) plants were grown in the greenhouse on soil (Fruhstorfer Erde, Hawita), and were watered with tap water and fertilized every other week with 1 ml/L WUXAL Super (Aglukon). Wheat was grown for 6 weeks, poplar for 8 months. Prior to chloroplast isolation, whole plants were incubated in darkness at room temperature for 2–3 days to minimize starch content. *Spinacia oleracea* (spinach) leaves were purchased from the local market and not incubated. Around 100-300 g of leaves were harvested per isolation and yielded around 1 ml of spinach or 200 µl of wheat or poplar cell-free extract.

### Percoll gradients for density centrifugation

Percoll step gradients were assembled by combining isotonic Percoll stock (90% v/v Percoll, 50 mM HEPES/KOH pH 8.0, 2 mM EDTA, 0.3 M Mannitol, 0.1% w/v BSA, 5% v/v ddH_2_O) and leaf homogenization buffer B (50 mM HEPES/KOH pH 8.0, 2 mM EDTA/NaOH pH 8.0, 0.3 M Mannitol, 5 mM β-mercaptoethanol). The specific steps of the gradients varied for each plant species, with gradients set at 80%/40%/20% (v/v) Percoll stock for wheat, 80%/40%/30% (v/v) Percoll stock for spinach, and 80%/50%/20% (v/v) Percoll stock for poplar. The individual gradient steps were carefully layered into a 50 ml conical centrifugation tube with a serological pipette, starting with the highest Percoll concentration. Per tube, 30 ml of solution was used (7 ml/12 ml/11 ml).

### Chloroplast isolation

Isolation and lysis procedure was adapted from Clark et al.^38^ All steps of the chloroplast isolation procedure were carried out at 4°C or on ice. Harvested plant material was divided into 50-100 g batches, leaves of wheat and poplar were cut into 3 cm stripes prior to processing in a blender (8011ES, Waring). The plant material was homogenized in buffer A (50 mM HEPES/KOH pH 8.0, 2 mM EDTA/NaOH pH 8.0, 0.3 M Mannitol, 0.1% (w/v) BSA, 0.6% PVP, 5 mM β-mercaptoethanol) in a pre-cooled bucket on high setting for three bursts of 5s, 5s and 2s, respectively. The ratio of tissue to buffer was kept between 1:3-6 (w/v), higher ratios were used when leaf material was more rigid. The homogenate was then filtered through four layers of sterile cloth (2 sheets of Miracloth and 2 layers of cheesecloth) into 250ml polycarbonate bottles by gentle hand pressure. Following filtration of the homogenate, chloroplasts were sedimented at 1000 g for 8 min. Pellets were resuspended in 5 ml homogenization buffer B (see above) and a maximum of 3.5 ml of the suspension was gently layered on top of the Percoll step gradient. Gradients were centrifuged at 10,000 g for 10 min at 4°C with the slowest acceleration and deceleration setting. Intact chloroplasts were collected at the interface of the two highest Percoll concentrations (see Fig. S6) with a serological pipette and washed with at least 3 volumes of washing buffer (50 mM HEPES/KOH pH 8.0, 2 mM EDTA/NaOH pH 8.0, 0.3 M Mannitol, 5 mM β-mercaptoethanol,) at 1000 g for 8 min. The washing step was repeated twice. The weight of the remaining pellet was measured and intact chloroplasts were gently resuspended in 1 ml lysis buffer (30 mM HEPES/KOH pH 7.7, 60 mM potassium acetate, 7 mM Magnesium acetate, 60 mM ammonium acetate, 5 mM DTT, 0.5 mM PMSF, 10 v/v glycerol) per gram of chloroplasts. The suspensions were flash frozen in liquid N_2_ and stored at -80°C for up to a month.

### Lysis of chloroplasts and preparation of S30 extract

Chloroplast S30 extracts were prepared on ice by thawing aliquots of isolated chloroplasts for 20 minutes, followed by gentle pipetting to resuspend the chloroplasts. Chloroplast envelopes were disrupted by repeatedly passing the suspension through a 0.5 mm (25G) x 40 mm needle (Braun Sterican REF 9186166) into a sterile syringe. During dispersion back into the 1.5 ml centrifuge tube, formation of a droplet at the end of the needle was induced through gentle push on the needle plunger (Fig. S6). Spinach chloroplasts underwent at least 15 passes, wheat and poplar chloroplasts underwent at least 30 passes. Subsequently, GTP and amino acid solutions were added from 1000x stocks to reach final concentrations of 40 μM of each amino acid and 0.1 mM GTP. The lysed chloroplasts were centrifuged at 30.000g for 30 minutes at 4°C, the supernatant was transferred to fresh tubes, cleared at 30.000g at 4°C for 30 minutes, transferred to fresh tubes and cleared again at 30.000g at 4°C for 20 minutes. The resulting supernatant was transferred to fresh tubes in aliquots, flash-frozen in liquid nitrogen and can be stored at -80°C for up to 2 years without loss of activity. Extracts can be refrozen after thawing using liquid nitrogen, although activity will lower over multiple freeze-thaw cycles. Effective chloroplast cell-free extracts exhibit a slight green color and typically demonstrate a protein concentration of at least 25 mg/ml (Supplementary Figure 6). These characteristics served as a preliminary indicator for troubleshooting prior to conducting CFE assays.

### Assembly and preparation of DNA templates

All constructs were cloned using the Golden Gate assembly method.^50^ Level 0 parts are compatible with the Marburg collection^51^ and adhere to the PhytoBrick standard.^52^ Level 0 parts were either amplified via PCR from genomic DNA of the respective plant, created via primer annealing and extension reactions or were taken from the MoChlo collection.^10^ Golden Gate reactions were performed in a total of 10µl volume. Level 0 parts were cloned using a BsmBI compatible entry vector (BBa_K2560002). Level 1 reactions were set up as follows: 20 fmol of plasmid DNA from each part, 10 fmol of an Amp/ColE1 backbone, 1 μl T4 DNA Ligase Buffer, 0.5 μl T4 DNA Ligase and 0.5 μl of BsaI-HFv2 (lvl1) or BsmBI-v2 (lvl0) restriction enzyme. Reactions were incubated in a thermocycler with 30 cycles of 37°C [BsmBI-v2: 42°C] (5 min) and 16°C (5 min) followed by a final digestion at 37°C [42°C] (10 min) and enzyme inactivation at 80°C (10 min). 5µl of Golden Gate reaction was used for chemical transformation of *E. coli*.

DNA was isolated from *E. coli* Top10 cells and purified employing either the NEB Monarch Plasmid Miniprep Kit or the Macherey-Nagel Nucleospin kit, following the protocols provided by the manufacturers. For experiments involving endogenous transcription, DNA was prepared in *E. coli* Epi400, chosen for its ability to adjust plasmid copy number and mitigate toxicity from potent chloroplast promoters. In such instances, plasmid copy number was increased with arabinose as per the manufacturer’s guidelines.

Plasmids containing endogenous chloroplast promoters were purified using Macherey-Nagel Nucleospin kit according to the manufacturer’s instructions and boiled for 30 minutes at 100°C to denature residual NanoLuc protein.

A list of all plasmids and sequences used in this study can be found in the Supporting Information (Supplementary Table S3).

### Preparation of translation buffer

Stock solutions were prepared in nuclease-free water. 2 M HEPES, 3.5 M KOAc, 3 M MgOAc, 2.9 M NH_4_OAc, 0.5 M ATP, 0.1 M GTP, 0.1 M CTP, 0.1 M UTP, 1 M creatine phosphate solutions and 50 mM each of 20 amino acids were titrated to pH 7.3 with KOH. 1 M DTT and 0.1 M spermidine were not pH titrated. Translation buffer was assembled on ice (15 mM HEPES, 60 mM KOAc, 10 mM MgOAc, 30 mM NH_4_OAc, 2 mM ATP, 1 mM GTP, 1 mM CTP, 1 mM UTP, 2 mM of 20 amino acids, 8 mM creatine phosphate, 5 mM DTT, 0.1 mM Spermidine), pH titrated to 7.3, frozen in liquid nitrogen and stored at -80°C.

### Cell-free reactions & NanoLuc assay

We assembled cell-free transcription and translation reactions from 50% extract, 20% DNA and 30% reaction buffer (consisting of 13% translation buffer and 17% other individual components) to yield final concentrations of 0.28 U/µl T7 RNA Polymerase, 0.025 U/µl Creatine Phosphokinase, 0.5 U/µl RNase inhibitor, 2% w/v PEG 3350, 1.95 mM HEPES pH 7.3, 7.86 mM KOAc, 1.31 mM MgOAc, 3.93 nM NH4OAc, 0.131 mM GTP, CTP, UTP, 0.262 mM ATP, 0.262 mM amino acids (each), 1.048 mM creatine phosphate, 0.655 mM DTT, 0.013 mM spermidine (see Tables S1 and S2 for an example). After at least 4 hours reaction time at RT, protein production was determined by endpoint measurements in a plate reader (Tecan Spark) using the Nano-Glo Luciferase Assay system (Promega REF N1110), by dispensing the Nano-Glo assay reagents at an equal volume to the protein synthesis reaction.

Reactions were manually prepared with a 10 µl reaction volume unless specified otherwise. Reactions prepared using liquid handling robots used 2 µl reaction volume unless otherwise stated. Reaction components were added using an Echo 525 liquid handling robot (Beckmann-Coulter). Liquid dispensing instructions were written using the PyEcho script (https://github.com/HN-lab/PyEcho). Nano-Glo assay reagents were dispensed using the Cobra Nano liquid handling robot (Art Robbins). Reaction vessels were either white 384 well plates (Corning REF 4513) covered with breathe-easy foil (Sigma Aldrich REF Z380059) or 1.5 ml reaction tubes.

### Data analysis and visualization

Data analysis and visualization were performed using Python 3.10.5. For parsing and processing, the pandas library (version 1.4.3) was utilized. Visualization of the data was conducted using the Plotly library (version 5.9.0). Linear least-squares regression analysis in figure 7 was performed on log-log transformed data using the SciPy library (version 1.8.1).

Data is displayed as box plots and adjacent individual data points on decadic logarithm scale. The midlines of the box plots represent the median, the boxes’ upper and lower limits represent the 1st and 3rd quartiles. Whiskers correspond to the box’ edges +/- 1.5 times the interquartile range.

### Calibration of luminescence output using NanoLuc

Purified NanoLuc protein (Promega REF G9711) was diluted to 10 µM in 0.1 mg/ml BSA solution. Cell-free reactions were set up manually in a total volume of 10 μl using 10μM UTC 7.0 DNA template. To account for the absorbance of the green cell-free extract during subsequent NanoLuc quantification, the samples were diluted 1:2 (v/v) with 0.1 mg/ml BSA solution prior to luminescence measurement and equal volumes of the cell-free extracts were added to the purified NanoLuc protein. Absolute NanoLuc concentrations in the reactions were calculated from a log10-transformed standard curve fitted to a line (Supplementary figure S3).

## Author Information

### Author Contributions

Clemens Böhm and René Inckemann contributed equally.

### Notes

The authors declare no competing financial interest.

## Acknowledgments

We thank the members of the iGEM Team Marburg 2021, who were involved in the ideation, planning and initial experiments of this project during the iGEM competition. We thank Arpita Sahoo and François-Xavier Lehr for the PyEcho script, Andrea Polle (Göttingen) for the generous provision of *Populus × canescens* trees, Ulrich Zick for the generous supply of *Triticum aestivum* seeds, and Christiane Rohrbach, Timo Engelsdorf and Julia Seufer for laboratory support and scientific guidance. This work was supported by the Max Planck Society. HN acknowledges funding by the Deutsche Forschungsgemeinschaft (DFG, German Research Foundation) grant NI 2040/1-1. MCJ and LC acknowledge funding by the U.S. Department of Energy (DE-SC0023278).

## Notes

### Competing Interest Statement

The authors have declared no competing interest.

