## Supplementary Information for "Chloroplast Cell-Free Systems from Different Plant Species as a Rapid Prototyping Platform"

### Corresponding Authors

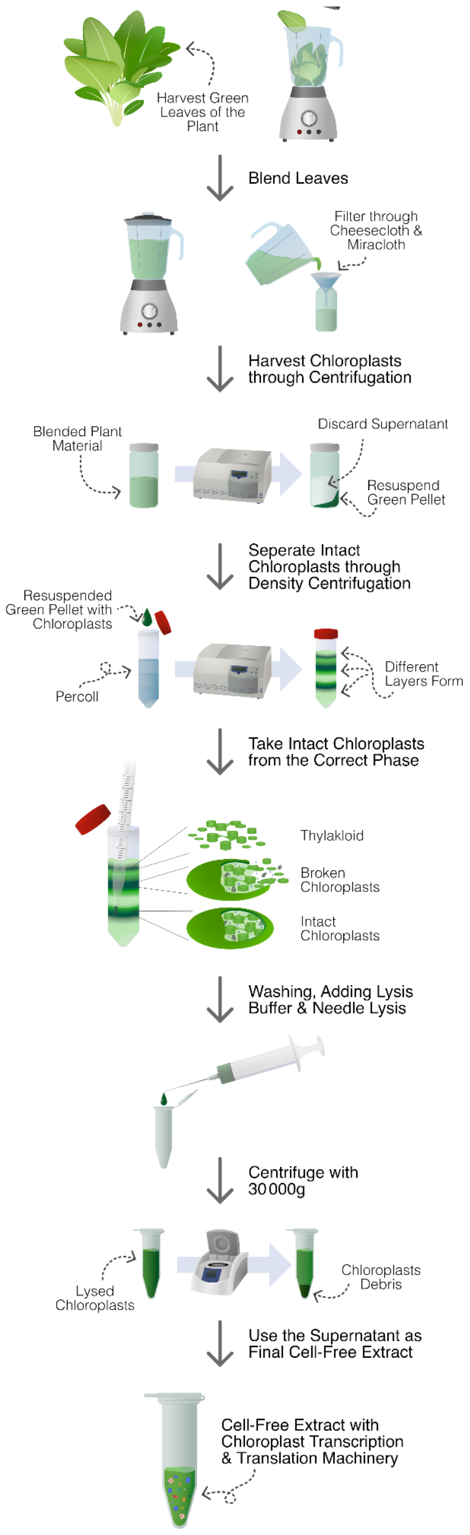


**Supplementary Figure S1**: **Workflow of chloroplast isolation.** Plant leaves were harvested and homogenized using a blender. After cloth filtering, the homogenate was centrifuged and the pellet resuspended. Percoll gradients were used to separate intact chloroplasts from broken ones. After washing, chloroplasts were lysed using a needle. Chloroplast debris was removed via ultracentrifugation and supernatant used as final cell-free extract.

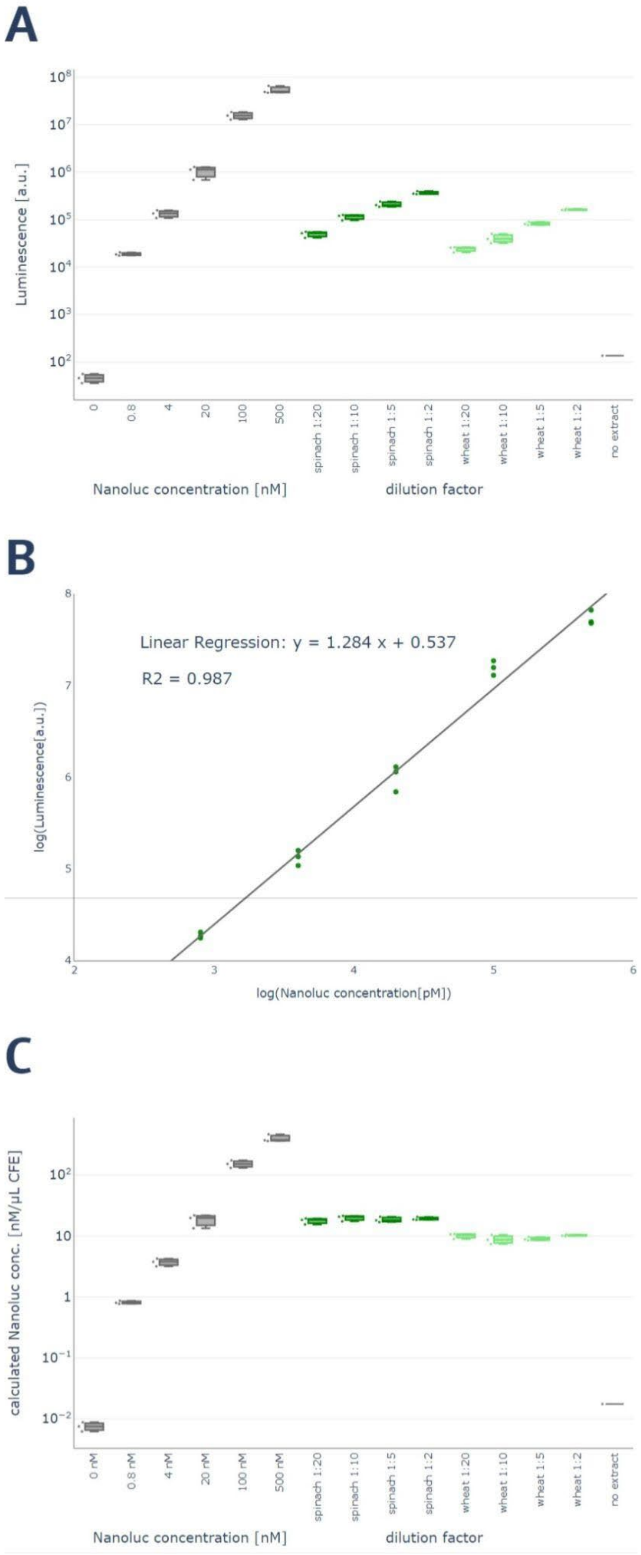


**Figure S2**: **Calibration of luminescence output of cell-free systems with purified NanoLuc.** (A) Raw luminescence values of purified NanoLuc (gray) and different dilutions of the spinach (dark green) and wheat (light green) cell-free systems after the synthesis reaction. (B) Calibration curve showing luminescence output as a function of NanoLuc protein concentration, including linear regression and $R^2$ value. (C) Calculated NanoLuc concentration per µl CFE reaction using data from the calibration curve in B, yielding 18.8 ± 1.8 nM for spinach and 9.5 ± 0.9 nM for wheat. Calculated total luminescence is very similar between dilution factors, indicating robust correction for absorption by the green extracts. Cell-free reactions were set up manually with a total volume of 10 µl and UTC 7.0 plasmid. Luminescence was measured after 4 hours of incubation at 20°C (N=3).

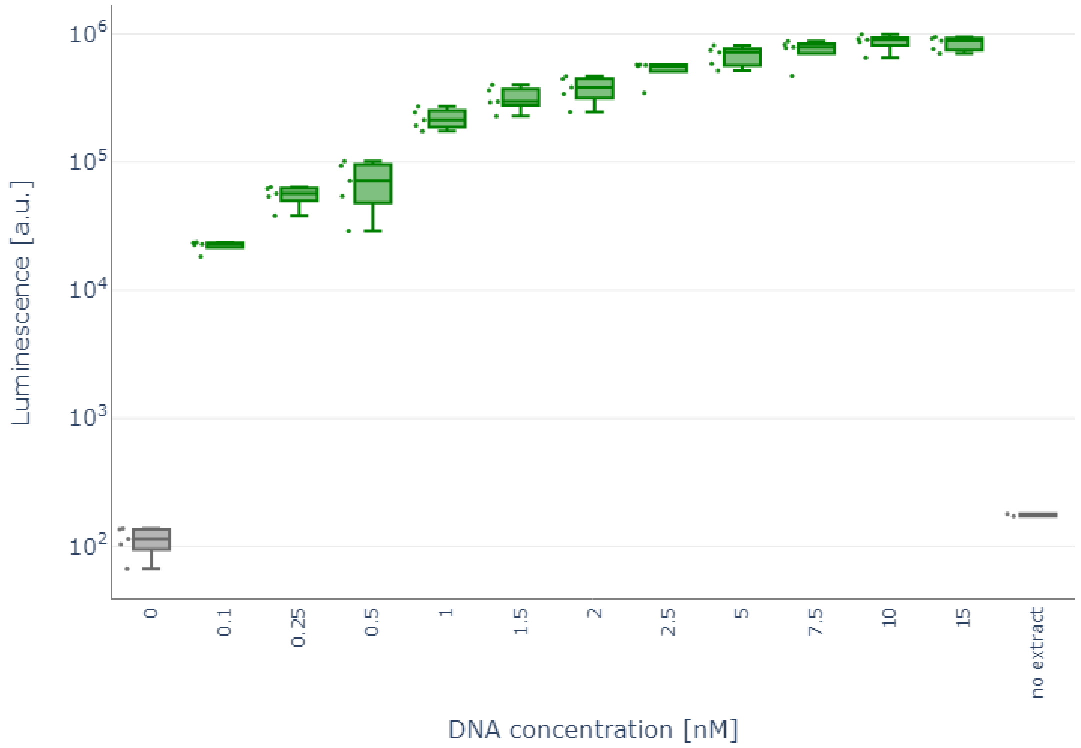


**Figure S3: Effect of DNA concentration on cell-free protein production.** NanoLuc luminescence signals obtained in CFE with varying template DNA concentrations. Highest expression is found at 10 nM UTC plasmid DNA concentration, all tested concentrations show expression above background. Negative controls either lack extract or DNA. Cell-free reactions were set up with a total volume of 2 µl and NanoLuc activity was measured after 4 hours of incubation at 20°C (N=5).

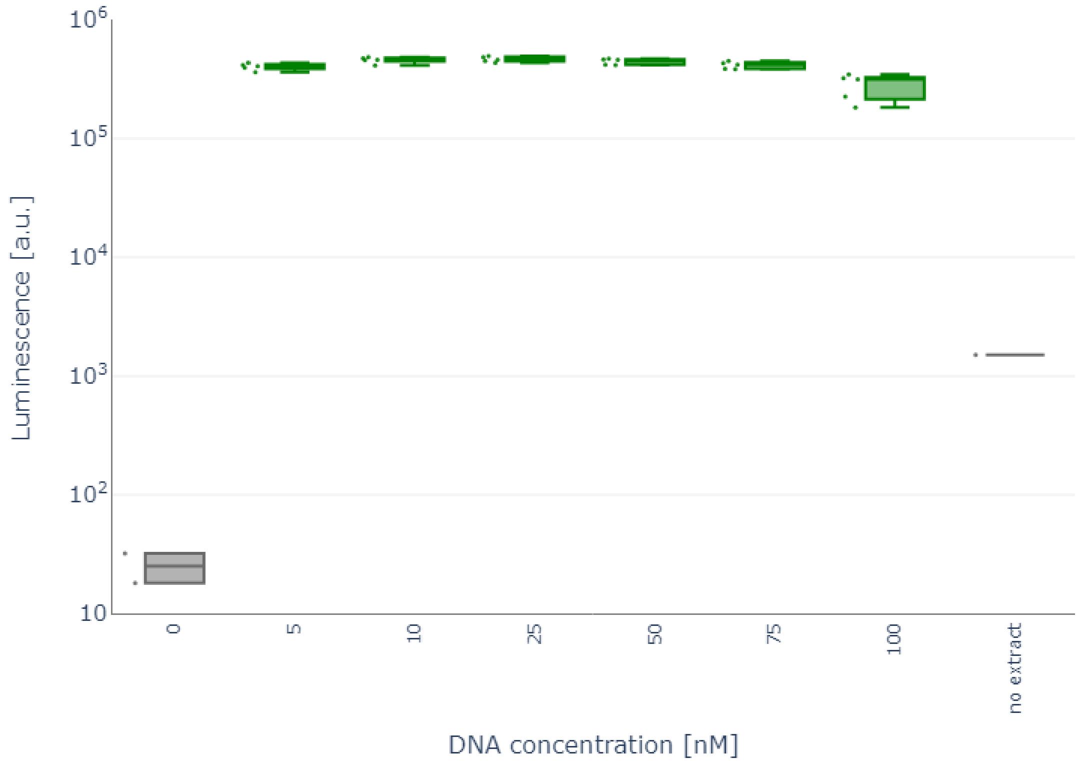


**Figure S4: Effect of high DNA concentration on cell-free protein production.** NanoLuc luminescence signals obtained in CFE with varying template DNA concentrations. Highest expression is found again at 10 nM UTC plasmid DNA concentration, all tested concentrations show expression above background. Negative controls either lack extract or DNA. Cell-free reactions were set up with a total volume of 2 µl and NanoLuc activity was measured after 4 hours of incubation at 20°C (N=5).

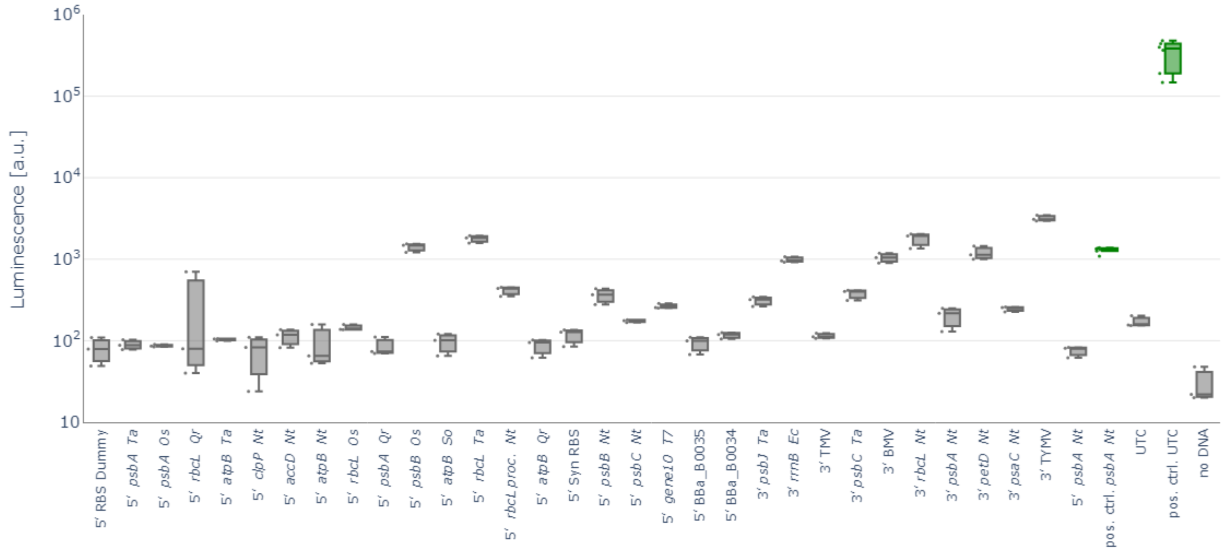


**Figure S5: Background luminescence of template DNA plasmids in absence of cell-free extract.** NanoLuc luminescence signals obtained in reactions lacking chloroplast extract. Negative control with included spinach extract lacks DNA. Positive controls containing all reagents were run alongside and measured at the same time. Mean expression was at least 17 times higher when extract was included. Reactions were set up with a total volume of 2 µl and NanoLuc activity was measured after 4 hours of incubation at 20°C (N=3).

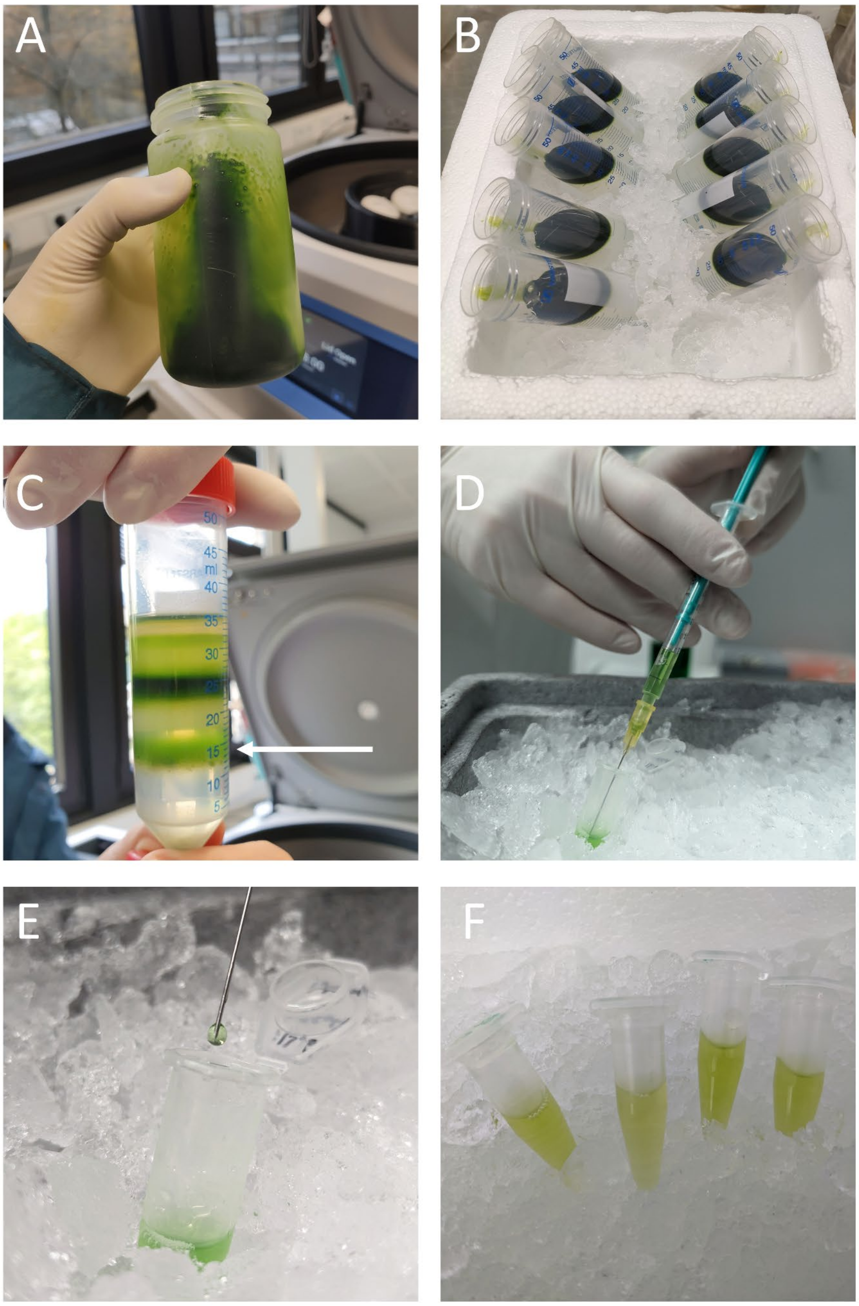


**Figure S6**: **Images of the chloroplast isolation workflow.** (A) Chloroplast pellet in centrifuge bottle after homogenization, filtering and first centrifugation. Importantly, very little starch is visible at this step. Starch, in high concentrations visible at this step as a white pellet, greatly reduces intact chloroplast yield. (B) Washed chloroplast suspension layered on top of Percoll step gradients. (C) Gradient after density centrifugation. White arrow indicates the position of intact chloroplasts. In layers above, broken chloroplasts and thylakoid membranes are visible. (D & E) Needle lysis of chloroplasts. During lysis (E), droplets are formed at the needle tip by applying gentle pressure on the needle plunger. (F) Final, functional cell-free extracts exhibit a slight green color.

### Supplementary Tables

**Table S1: Translation buffer components and stock solutions.**

| Reagent for translation buffer | Stock concentration [mM] | Final concentration in translation buffer [mM] | Stock volume per µl reaction [µl] |
|---|---|---|---|
| HEPES | 2000 | 15 | 0.008 |
| KOAc | 3500 | 60 | 0.017 |
| MgOAc | 3000 | 10 | 0.003 |
| $NH_4OAc$ | 2900 | 30 | 0.01 |
| ATP | 500 | 2 | 0.004 |
| GTP | 100 | 1 | 0.01 |
| CTP | 100 | 1 | 0.01 |
| UTP | 100 | 1 | 0.01 |
| Creatine Phosphate | 1000 | 8 | 0.04 |
| 20 amino acids (each) | 50 | 2 | 0.008 |
| DTT | 1000 | 5 | 0.005 |
| Spermidine | 100 | 0.1 | 0.001 |

**Table S2: Reaction components and stock solutions.**

| Reagent for reaction mastermix | Stock concentration | Final concentration in CFE reaction | Stock volume per µl reaction [µl] |
|---|---|---|---|
| Translation buffer | - | 13% v/v | 0.13 |
| Creatine kinase | 7.5 U/µl | 0.025 U/µl | 0.033 |
| T7 Polymerase | 20 U/µl | 0.28 U/µl | 0.014 |
| PEG 3350 | 20% w/v | 2% w/v | 0.1 |
| RNase Inhibitor | 40 U/µl | 0.5 U/µl | 0.013 |
| Plasmid Template | 50 nM | 10 nM | 0.2 |

**Table S3: Table of all regulatory elements used in this study.**

| Type | Name | Sequence | length (bp) | Description |
|---|---|---|---|---|
| Promoter | T7 promoter | TAATACGACTCACTATAG | 18 | core promoter region of the T7 promoter from *Escherichia* phage T7 |
| Promoter | P*rrn16* Nt | GCTCCCCCGCCGTCGTTCAATG AGAATGGATAAGAGGCTCGTGG GATTGACGTGAGGGGGCAGGGA TGGCTATATTTCTGGGAGCGAAC TCCGGGCGAATACGAAGCGCTT GGATACAGTTGTAGGGAGGGAT TT | 135 | promoter of the plastidial 16S ribosomal RNA (*NitaCr105*) of *Nicotiana tabacum* |
| Promoter | P*rrn16* Cr | CAGGCAACAAATTTATTTATTGTC CCGTAAGGGGAAGGGGAAAACA ATTATTATTTTACTGCGGAGCAG CTTGTTATTAGAAATTTTTATTAA AAAAAAAATAAAAATTTGACAAAA AAAAATAAAAAAGTTAAATTAAAA ACACTGGGAATGTTCTAACAATC ATAAAAAAATCAAAAGGGTTTAA AATCCCGACAAAATTTAAACTTTA AAGAGT | 217 | promoter of the plastidial 16S ribosomal RNA (*CreCp.r003100.rRNA*) of *Chlamydomonas reinhardtii* |
| Promoter | P*rrn16* Ta | ATAAGAGGCTTGTGGGATTGAC GTGATAGGGTAGGGTTGGCTAT ACTGCTGGTGGCGAACTCCAGG CTA | 69 | promoter of the plastidial 16S ribosomal RNA (*TraeCr087*) of *Triticum aestivum* |
| Promoter | P*psbA* Nt | GATCTACATACACCTTGGTTGAC ACGAGTATATAAGTCATGTTATA CTGTTG | 52 | promoter of Photosystem II protein D1 (*psbA; NitaCp001)* of *Nicotiana tabacum* |
| Promoter | P*rbcL* Nt | GGGGGAAGTTCTTATTATTTAGG TTAGTCAGGTATTTCCATTTCAAA AAAAAAAAAAGTAAAAAAGAAAA ATTGGGTTGCGCTATATATATGA AAGAGTATACAATAATG | 110 | promoter of Ribulose bisphosphate carboxylase large subunit (*rbcL*; *NitaCp031*) of *Nicotiana tabacum* |
| 5'UTR | *accD* 5'UTR Nt | AAGTGTTCCCCCAGATTCAGAAC TTTTTTTCAATACTCACAATCCTT ATTAGTTAATAATCCTAGTGATTG GATTTCTATGCTTAGTCTGATAG GAAATAAGATATTCAAATAAATAA | 180 | 5'untranslated region of acetyl-CoA carboxylase beta subunit (*accD*; *NitaCp032*) |

| | | | | |
|---|---|---|---|---|
| | | TTTTATAGCGAATGACTATTCATC TATTGTATTTTCATGCAAATAGG GGGCAAGAAAACTCT | | of *Nicotiana tabacum* |
| 5'UTR | *atpB* 5'UTR Nt | CAAATGAAAGACTTTCTCAAGAT TCTGATTCATCCACTTGAGATTTT AAAATTAAAATAGGTTGGGTGGG CTTGCAAATTCACTCAGTCTCAG TGAATAAGTAAACAATTGAATCG GTTCAATTGCATGGTGCCAACGA AATCGAGTGCTAATTCCCATTTT ATTGAATTAACCGATCGACGTGC TAGCGGACATTTATTTTGAATTC GATAATTTTTGCAAAAACATTTCG ACATATTTATTTATTTTATTATT | 255 | 5'untranslated region of ATP synthase subunit beta (*atpB*; *NitaCp030*) of *Nicotiana tabacum* |
| 5'UTR | *atpB* 5'UTR Qr | CCTAGATGTGAAAATAGGAGGA GTTCGCCCATGAAAAGCAAAGCA TGAAACTAAAACTCTAAAACTAAA GGGTATAGGTAAAAAAAAAATAG GCTAGGCATAAATCGATAGGCTT AAATATTAACTAAGAAATGAGATA AGGGCACCAATAAGATAGAAAAA ATGAATCGTAAATAGAAATAGAG TTCCGGTTCGAATTCGATAAATA ATATGGATGATATTGTCTATAATG ATAGTCAAATGAAAGACTTTCTC AAGACTTTTATTGATCCGCCTGA GATTTTGAAAATGAGTTGGTTGA ACTTGAAAATTAACTCATTGAAAT TGAATAAATAAACAATCGAATTG GATTCGATTGGATGGTACCAACG AAATCTAGTGCAGTGCGAAACCC CATTTATTATGGAATTATTATTGA ATTAACCGATCAACTTGCTTTTCT ATCGAACATTTTTTTTATTTCAAT AATTTTCGCAAAAAAAAATTTCGA CATATTATATTTTATT | 506 | 5'untranslated region of ATP subunit beta (*atpB*; *HCS81_pgp061*) of *Quercus robur* |
| 5'UTR | *atpB* 5'UTR So | GTGAAAATATGCAGAATTCTCTC ATGAAAGGATAAAAGAATAGGCT ACTCATAAATCTATATACTAAATC GAAACTAAGTCCCAGTACGATAG AAATAATGAATCATAAAAAAATAT AGTTTTAGAGTTCGGGTTCGATT TCCATAGATAATCTAGAAAGGAG TGTCTATAATGATAGGCAAATAA AAGACTTTCTCGGGATTTTTGGT CATCCGTTTGATATTTTGAAAATA GGCGGATTGCAATTTCAAATTGA ATAGAAATAGAATAATTCAATTCC AAAAAGTAAACAATTGAATTGGA GTCCTTTTTTTTTGCTGGTACCAA CAAAATTTATTGCTAACCCCTATT | 453 | 5'untranslated region of ATP subunit beta (*atpB*; *SpolCp032*) of *Spinacia oleracea* |

| | | | | |
|---|---|---|---|---|
| | | TCTTATTTAATTAATCGATCAGCT TGCTATCGGACATTTTTTTTATTT TGGATTCGATAATTTTCATTTTTG CAAAAAATTTCGACATACTTTACT ATATATT | | |
| 5'UTR | *atpB* 5'UTR Ta | TTTGTATATCGAAGTCCTAGATA GGAAAGTAGAGTAGGCACAGAT CCTCCACAAAAGGCAAAATGTAT ATGAAAAAAAGATTGATTGAACT TTCCAACGGACTCATTCCATGAG TAAACGATTGAATGGGATTCGCT TGGGCAACGAAATCAAGTCCTG GTCCCCTTTTCTCTCTTATTGAAT TAACTAATTCATTTCCTTTTTACT TTTGGATTTTTTTTGATTTGATTT GGCATTATTCAACAATAAAAAAA GAAAAATTTCGACAAATTCCTTTT TTTTAATTATGTGATAATT | 297 | 5'untranslated region of ATP subunit beta (*atpB*; *TraeCp029*) of *Triticum aestivum* |
| 5'UTR | *clpP* 5'UTR Nt | GTTTCCACCTCAAAGTGAAATAT AGTATTTAGTTCTTTCTTTCATTT A | 48 | 5'untranslated region of ATP-dependent Clp protease proteolytic subunit (*clpP*; ) of *Nicotiana tabacum* |
| 5'UTR | *gene10* 5'UTR T7 | GGCAGACCACAACGGTTTCCCA CTAGAAATAATTTTGTTTAACTTT AAGAAGGAGATATACAT | 63 | 5'untranslated region of Major capsid protein (*gene10*; *T7p45*) of *Escherichia* phage T7 |
| 5'UTR | *psbA* 5'UTR Nt | AATAAAAAGCCTTCCATTTTCTAT TTTGATTTGTAGAAAACTAGTGT GCTTGGGAGTCCCTGATGATTAA ATAAACCAAGATTTTACC | 88 | 5'untranslated region of Photosystem II protein D1 (*psbA*; *NitaCp001*) of *Nicotiana tabacum* |
| 5'UTR | *psbA* 5'UTR Os | TAACAAGCCTTCTATTATCTTTCT AGTTAATACGTGTGCTTGGGAGT CCTTGCAATTTGAATAAACCAAG ATCTTACC | 78 | 5'untranslated region of Photosystem II protein D1 (*psbA*; *AKK66_gp001*) of *Oryza sativa* |
| 5'UTR | *psbA* 5'UTR Qr | TAACAAGCCCTCAATTATCTATTT CTATTTATAGAGAATCGTGTGCT TGGGAGTCCCTGATGATTAATGA TTAAATAAACCAAGATTTTACC | 92 | 5'untranslated region of Photosystem II protein D1 (*psbA*; *HCS81_pgp089*) |

| | | | | |
|---|---|---|---|---|
| | | | | of *Quercus robur* |
| 5’UTR | *psbA* 5’UTR So | TAACAATCTTTCAATTTCTATTTC TAGCGAATTTGTGTGCTTGGGAG TCCCTGATGATTAAATTAAATAAA CCAAGATTTTACC | 84 | 5’untranslated region of Photosystem II protein D1 (*psbA*; *SpolCp002*) of *Spinacia oleracea* |
| 5’UTR | *psbA* 5’UTR Ta | TAACAAGCCTCCTATTATCTATAT TCTAGTTAATACGTGTGCTTGGG AGTCCTTGCAATTTGAATAAACC AAGATCTTACC | 81 | 5’untranslated region of Photosystem II protein D1 (*psbA*; *TraeCp001*) of *Triticum aestivum* |
| 5’UTR | *psbB* 5’UTR Nt | GCTTCTCTTTGTTCCTACGAACA GAATTGTTCCATTATTACCAACA GAATAGAACACCCTTGTTCGGAA ATAATCGACTGAACAAGAGTGGT CCATAGGATAGTCATATTATAGT CTTTTCCAATGCAATAAAGTTAC GTAGTGTCTATTTATCTTTGATAT AAGGGGTATTTCC | 175 | 5’untranslated region of Photosystem II CP47 reaction center protein (*psbB*; *NitaCp052*) of *Nicotiana tabacum* |
| 5’UTR | *psbB* 5’UTR Os | GTATAGAATAGATCTGCTTCTCT TTCTTCTTACGAACAGAATTGGC TTCTTATTTTTAATGGAATGAAAT AAATATTCACGCTTTCTGACACA GAATCCCCTAGAAGGGTTAGGTA CATAGGATATGGATAGTCTTTGC CAATGCGATAAAATAAAGTGACA TCGTGTCTATTTTTCTTTGCTAAA GGGGTATTTCC | 197 | 5’untranslated region of Photosystem II CP47 reaction center protein (*psbB*; *AKK66_gp055*) of *Oryza sativa* |
| 5’UTR | *psbC* 5’UTR Nt | GTTATTTGTACCAGTAACCGGTT TATGGATGAGTGCTCTTGGAGTA GTCGGTCTAGCCCTGAACCTAC GTGCCTATGACTTCGTTTCTCAG GAAATTCGCGCAGCGGAAGATC CTGAATTTGAGACTTTCTACACC AAAAATATTCTCTTAAACGAAGG TATTCGCGCTTGGATGGCGGCT CAAGATCAGCCTCATGAAAACCT TATATTCCCTGAGGAGGTTCTAC CAC | 230 | 5’untranslated region of Photosystem II CP43 reaction center protein (*psbC*; *NitaCp016*) of *Nicotiana tabacum* |
| 5’UTR | *rbcL* 5’UTR Os | GGATTTGGTGAATCAAATCCATG GTTTAATAACGAAGCATGTTAAC TTACCATAACAACAACTCAATTCT TATCGAATTCCTATAGTAGAATTC CTATAGCATAGAATGTACACAGG GTGTACCCATTATATATGAATGA AACATATTATATGAATGAAACATA | 325 | 5’untranslated region of Ribulose bisphosphate carboxylase large subunit (*rbcL*; *AKK66_gp073*) of *Oryza sativa* |

| | | | | |
|---|---|---|---|---|
| | | TTCATTAACTTAAGCATGCCCCC CATTTTCTTTAATGAGTTGATATT AATTGAATATCTTTTTTTTAAGAT TTTTGCAAAGGTTTCATTTACGC CTAATCCATATCGAGTAGACCCT GTCGTTGTGAGAATTCTTAATTC ATGAGTTGTAGGGAGGGACGT | | |
| 5'UTR | *rbcL* 5'UTR Qr | GTATTTGGCGAATCAAATATCAT GGTCTAATAACGAACCATTCTAA TTAGTTGATAATTTTTTGAAAGGA TTCCTTGAAAGGTTTCATTAACTC CTAATTCATGTCGAGTAGACCTT GTTGTTGCGAAAATTCTTAATTC | 140 | 5'untranslated region of Ribulose bisphosphate carboxylase large subunit (*rbcL*; *HCS81_pgp060*) of *Quercus robur* |
| 5'UTR | *rbcL* 5'UTR Ta | GGATTTGGTAAATCAAATCCATG GTTTAATAACGAACCGTGTTAAC TTACCATAACAACAACTCAATTC CTATCGAATTCCTATAGTGGAAT TCCTATAGGATAGAACATACACA GGGTGTACGCATTATATATGAAT GAAACATATTCATTAACCTAAGC ATGCCCTCAATTTTCTTTAATGAG TTGATATTATATTAATTGAATATC CTTTTTGTTTTACGAGATTTTTGC TAAAGTTTCATTTACGCCTAATTA ACATCGAGTAGACCCTGTTATTG TGAGAATTCTTAATTCAAGAGTTA TAGGGAGGGACTT | 317 | 5'untranslated region of Ribulose bisphosphate carboxylase large subunit (*rbcL*; *TraeCp030*) of *Triticum aestivum* |
| 5'UTR | *rbcL* 5'UTR So | AAATACATGGTCTATTAACGAAC CATTTTGATTAGTTGATAATATTA ATTGAGAATTTGATGAAAGATTG CTATAAAAGGTTTCATTAAGGCC TAATTTATGTCGAGTAGACCTTG TTGCTTTGTTGTAAAAATTAAAAT TTGAAGTTGTAGGGAGGGACTT | 176 | 5'untranslated region of Ribulose bisphosphate carboxylase large subunit (*rbcL*; *SpolCp033*) of *Spinacia oleracea* |
| 5'UTR | *rbcL* 5'UTR processed Nt | GTCGAGTAGACCTTGTTGTTGTG AGAATTCTTAATTCATGAGTTGTA GGGAGGGATTT | 58 | processed version of 5'untranslated region of Ribulose bisphosphate carboxylase large subunit (*rbcL*; *NitaCp031*) of *Nicotiana tabacum* |
| RBS | BBa_B0034 | AGAGAAAGAGGAGAAATAATC | 21 | RBS B0034 of the Community RBS part collection |
| RBS | BBa_B0035 | AGAGATTAAAGAGGAGAATAATC | 23 | RBS B0035 of the Community RBS |

| | | | | |
|---|---|---|---|---|
| | | | | part collection |
| RBS | Synthetic RBS | AAATTCGATAGAGATGAAATTGGAGCTCTAGAGAATTCAGTTGTAGGGAGGGATCC | 56 | Synthetic RBS based on the 5'untranslated region of *rbcL* (*NitaCp031*) of *Nicotiana tabacum* |
| RBS | RBS_Dummy | AGAGTGTCAGGATACCCGATAATC | 24 | 5'untranslated region with low translational activity |
| CDS | NanoLuc | GTTTTCACTTTAGAAGACTTCGTAGGTGACTGGCGTCAAACAGCAGGTTATAATTTAGACCAAGTTTTAGAACAAGGTGGTGTATCAAGTTTATTCCAAAATTTAGGTGTTTCTGTTACTCCAATTCAACGTATCGTATTAAGTGGTGAAAATGGTCTTAAAATTGACATCCATGTTATTATTCCTTATGAAGGTCTTTCAGGTGACCAAATGGGTCAAATTGAAAAAATTTTTAAAGTAGTTTATCCTGTAGATGACCATCACTTTAAAGTAATTTTACACTATGGTACTTTAGTAATTGATGGCGTTACACCTAATATGATTGACTACTTTGGTCGTCCTTATGAAGGTATTGCTGTTTTTGATGGTAAAAAAATCACAGTTACAGGTACATTATGGAATGGTAATAAAATTATTGACGAACGTTTAATCAATCCTGATGGTTCATTATTATTCCGTGTTACAATTAATGGTGTTACAGGTTGGCGTCTTTGTGAACGTATCCTTGCT | 510 | Coding sequence of the NanoLuc luciferase |
| 3'UTR | *petD* 3'UTR Nt | AAATTTTTAAAGATTCAATTGTGAAATAACACGACATGTGTATCTAGGGAATAGTTTCTTCAAAGCGAATTCTCCCTAGATACATCTATTCAATTTAATTCTGAATTTATTTTGAATATATGATATATTAATATATTAATTGTGCTAAAGAGTTTCAATCTATTTTCACTAAGTAAGTCCAATAGAT | 187 | 3'untranslated region of Cytochrome b6-f complex subunit 4 (*petD*; *NitaCp051*) of *Nicotiana tabacum* |
| 3'UTR | *rpoA* 3'UTR Nt | AAATCTATTGGACTTACTTAGTGAAAATAGATTGAAACTCTTTAGCACAATTAATATATTAATATATCATATATTCAAAATAAATTCAGAATTAAATTGAATAGATGTATCTAGGGAGAATTCGCTTTGAAGAAACTATTCCCTAGATACACATGTCGTGTTATTTCACAATTGAATCAATTTAAAA | 189 | 3'untranslated region of DNA-directed RNA polymerase subunit alpha (*rpoA*; *NitaCp057*) of *Nicotiana tabacum* |

| | | | | |
|---|---|---|---|---|
| | | AT | | |
| 3'UTR | *psaC* 3'UTR Nt | AATGATACGTTCGAGAAAACTCTACTTGAATCCATTTAATTTTTTTACCGACAAACCTGTGCTCGAAAATCACAATATTTTGAGCACGGGTTTTTATG | 99 | 3'untranslated region of Photosystem I iron-sulfur center (*psaC*; *NitaCp084*) of *Nicotiana tabacum* |
| 3'UTR | *psbA* 3'UTR Nt | AATCCTGGCCTAGTCTATAGGAGGTTTTGAAAAGAAAGGAGCAATAATCATTTTCTTGTTCTATCAAGAGGGTGCTATTGCTCCTTTCTTTTTTT | 95 | 3'untranslated region of Photosystem II protein D1 (*psbA*; *NitaCp001*) of *Nicotiana tabacum* |
| 3'UTR | *psbC* 3'UTR Ta | AAGATTTTCTTATTTATACCTGTTCTACTTTTTTTCTGTTCTGGCTCGGTTATTCCATCTAGCCGAGCCATTCATTCCTTTTTATGAAAGAAAGATAAGGGACAGAAAAAAAAAAAAAA | 119 | 3'untranslated region of Photosystem II CP43 reaction center protein (*psbC*; *TraeCp007*) of *Triticum aestivum* |
| 3'UTR | *psbJ* 3'UTR Ta | AATAATCGGAGGGACCAGATTGTAAACATGAAAAAGTAGGAGCTTAGCGGGTCCTTACCCCCCTTTATCTGATTAGAGCGGAAAGGACCCGCGGAATTTTTACTCTTATAACGCGAATTGATTCTATTCGATTCACTCTTATGAAGCAACAAGAAAAAGAGATCACTCGAGGATCCAATATCTTATTCCACGAAGGAAGTATCCTGGAAATCCTTGATTTAGTTTCGAGTAATAAACTAATAAAACCTTAATCAAAACTATTCAACTAGCCTAAAAAATAAAAAAAAA | 288 | 3'untranslated region of Photosystem II reaction center protein J (*psbJ*; *TraeCp036*) of *Triticum aestivum* |
| 3'UTR | *rbcL* 3'UTR Nt | AAAAACAGTAGACATTAGCAGATAAATTAGCAGGAAATAAAGAAGGATAAGGAGAAAGAACTCAAGTAATTATCCTTCGTTCTCTTAATTGAATTGCAATTAAACTCGGCCCAATCTTTTACTAAAAGGATTGAGCCGAATA | 142 | 3'untranslated region of Ribulose bisphosphate carboxylase large subunit (*rbcL*; *NitaCp031*) of *Nicotiana tabacum* |
| 3'UTR | *rrnB* 3'UTR Ec | AAGTAGAAACGCAAAAAGGCCATCCGTCAGGATGGCCTTCTGCTTAATTTGATGCCTGGCAGTTTATGGCGGGCGTCCTGCCCGCCACCCTCCGGGCCGTTGCTTCGCAACGTTCAAATCCGCTCCCGGCGGATTT | 222 | 3'untranslated region of the ribosomal RNA operon copy B (*rrnB*; *b3971*) of *Escherichia coli* |

| | | | | |
|---|---|---|---|---|
| | | GTCCTACTCAGGAGAGCGTTCA CCGACAAACAACAGATAAAACGA AAGGCCCAGTCTTTCGACTGAG CCTTTCGTTTTATTTGATG | | |
| 3’UTR | TMV 3’UTR | AAATAATAAATAACGGATTGTGT CCGTAATCACACGTGGTGCGTA CGATAACGCATAGTGTTTTTCCC TCCACTTAAATCGAAGGGTTGTG TCTTGGATCGCGCGGGTCAAAT GTATATGGTTCATATACATCCGC AGGCACGTAATAAAGCGAGGGG TTCGAATCCCCCCGTTACCCCCG GTAGGGGCCCA | 192 | 3’untranslated region of Capsid protein (*TMVgp6*) of Tobacco Mosaic Virus |
| 3’UTR | TYMV 3’UTR | AAGTTCTCGATCTTTAAAATCGTT AGCTCGCCAGTTAGCGAGGTCT GTCCCCACACGACAGATAATCG GGTGCAACTCCCGCCCCTTTTCC GAGGGTCATCGGAACC | 107 | 3’untranslated region of Coat protein (*TYMVgp3*) |
| 3’UTR | BMV 3’UTR | AAGGTGCCTTTGAGAGTCTACTT TTGCTCTCTTCGGAAGAACCCTT AGGGGTTCGTGCATGGGCTTGC ATAGCAAGTCTAGATGCGGGTAC CGTACAGTGTTGAAAAACACTGT AAATCTCTAAAAGAAACCA | 133 | conserved 3’untranslated region of all Brome Mosaic Virus RNAs (1,2 and 3) |
